# Synergistic RAS-MAPK and AKT Activation in MYC-Driven Tumors via Adjacent PVT1 Rearrangements

**DOI:** 10.1101/2025.02.17.638454

**Authors:** Ashutosh Tiwari, Utkarsha Paithane, Jordan Friedlein, Kojiro Tashiro, Olivier Saulnier, Karina Barbosa, Quang Trinh, Bryan Hall, Shrawantee Saha, Aditi Soni, Takuma Nakashima, Andrey Bobkov, Lynn Miya Fujimoto, Rabi Murad, Svetlana Maurya, Mayank Saraswat, Shahab Sarmashghi, Joshua T. Lange, Sihan Wu, Meher Beigi Masihi, Srija Ghosh, Gazal Hemmati, Owen Chapman, Liam Hendrikse, Brian James, Jens Luebeck, Tanja Eisemann, Theophilos Tzaridis, Deepak Rohila, Robyn Leary, Jyotika Varshney, Badrinath Konety, Scott M. Dehm, Yasuhiko Kawakami, Rameen Beroukhim, David A Largaespada, Lincoln Stein, Lukas Chavez, Hiromichi Suzuki, William A Weiss, Jianhua Zhao, Aniruddha Deshpande, Robert J. Wechsler-Reya, Michael D. Taylor, Anindya Bagchi

## Abstract

MYC-driven (MYC+) cancers are aggressive and often fatal. MYC dysregulation is a key event in these cancers, but overexpression of MYC alone is not always enough to cause cancer. *Plasmocytoma Variant Translocation 1* (*PVT1*), a long non-coding RNA (lncRNA) adjacent to MYC on chromosome 8 is a rearrangement hotspot in many MYC+ cancers. In addition to being co-amplified with MYC, the genomic rearrangement at PVT1 involves translocation, which has had obscure functional consequences. We report that translocation at the PVT1 locus cause asymmetric enrichment of 5’-PVT1 and loss of 3’-PVT1. Despite being classified as a non-coding RNA, the retained 5’ region of PVT1 generates a circular RNA (CircPVT1) that codes for the novel peptide we call Firefox (FFX). FFX augments AKT signaling and synergistically activates MYC and mTORC1 in these cells. Further, the 3’ end of PVT1, which is lost during the translocation, codes for a tumor-suppressing micropeptide we named as Honeybadger (HNB). We demonstrate that HNB interacts with KRAS and disrupts the activation of KRAS effectors. Loss of HNB leads to activation of RAS/MAPK signaling pathway, and enhances MYC stability by promoting phosphorylation of MYC at Ser^62^. These findings identify PVT1 as a critical node that synchronizes MYC, AKT, and RAS-MAPK activities in cancer. Our study thus identifies a key mechanism by which rearrangements at the PVT1 locus activate additional oncogenic pathways that synergize with MYC to exacerbate the aggressiveness of MYC+ cancers. This newfound understanding explains the poor prognosis associated with MYC+ cancers and offers potential therapeutic targets that could be leveraged in treatment strategies for these cancers.

## INTRODUCTION

Many cancers have a copy number gain in the 8q24 region ^1 2 3 4 5 6 7^. This genomic amplification event has traditionally been thought to promote dysregulation of the MYC oncogene, but this has been based on the assumption that *MYC* is the sole causal gene amplified in this region. However, we have previously shown that in 98% of MYC+ cancers, *MYC* is co-amplified with *PVT1*, a long non-coding RNA (lncRNA) gene located ∼30kb downstream of *MYC* ^8^. Importantly, MYC is critically dependent on PVT1 levels: inhibition of PVT1 results in reduced MYC protein levels and impaired tumor growth, indicating that PVT1 plays a critical role in MYC-driven tumorigenesis ^8^. The *PVT1* locus, in addition to being commonly co-amplified with MYC, is also a hotspot for chromosomal translocations ^9 10 11 12 13^. These rearrangements are strongly linked to poor outcomes in cancer patients, suggesting their role in driving the aggressiveness of MYC+ cancers ^14 15^. Based on these observations, we hypothesized that PVT1 genomic rearrangements, including amplifications and translocations, significantly contribute to the aggressive nature of MYC+ cancers. Here we illuminate the mechanisms underlying the deleterious consequences of the rearrangement events at the *PVT1* locus by conducting in-depth functional analyses of the rearrangement of the *PVT1* locus. Our results identify MYC/PVT1 amplification and PVT1 translocation as key drivers of cancer pathogenesis, reshaping the oncogenic signaling landscape through the expression—or lack thereof—of previously uncharacterized short proteins.

## RESULTS

### Genomic rearrangements in PVT1 are prevalent in human cancers and result in asymmetric enrichment of the 5’-PVT1 region

To comprehensively understand the prevalence and nature of translocations at the PVT1 locus, we analyzed gene fusion data from the Cancer Cell Line Encyclopedia (CCLE) dataset^16^ (Figure 1A). We found that gene fusions involving PVT1 were the most common in the entire dataset, (Figure 1B, Supplementary Table 1), comprising both intrachromosomal (56%) and interchromosomal (44%) fusion events (Figure 1C, D, Supplementary Table 2, 3). Unlike other recurrent gene fusions, such as EWS-FLI1, BCR-ABL, and EML4-ALK, which are preferentially enriched in specific cancer subtypes, PVT1 gene fusions occurred across cancer types (Supplementary Figure 1A, Supplementary Table 4). This suggests a possible broader role of PVT1 translocation as a nodal locus in tumorigenesis. We also noted that the majority of PVT1 gene fusions were not frame-retaining, which means that they did not result in the formation of chimeric oncoproteins. This suggests that PVT1 rearrangements may enhance oncogenicity through other mechanisms. A more detailed analysis of the rearrangements revealed that 88% of fusion transcripts retained the 5’-PVT1 region, while only 12% retained the 3’-PVT1 region (Figure 1E, Supplementary Table 5). Examination of the gene structures resulting from *PVT1* rearrangements/translocations revealed that most of the intrachromosomal fusion events originated from extrachromosomal circular DNA (ecDNA) or chromosomally mediated mechanisms forming homogenously staining regions, (HSRs) (Supplementary Figure 1B-E).

**Figure 1:**
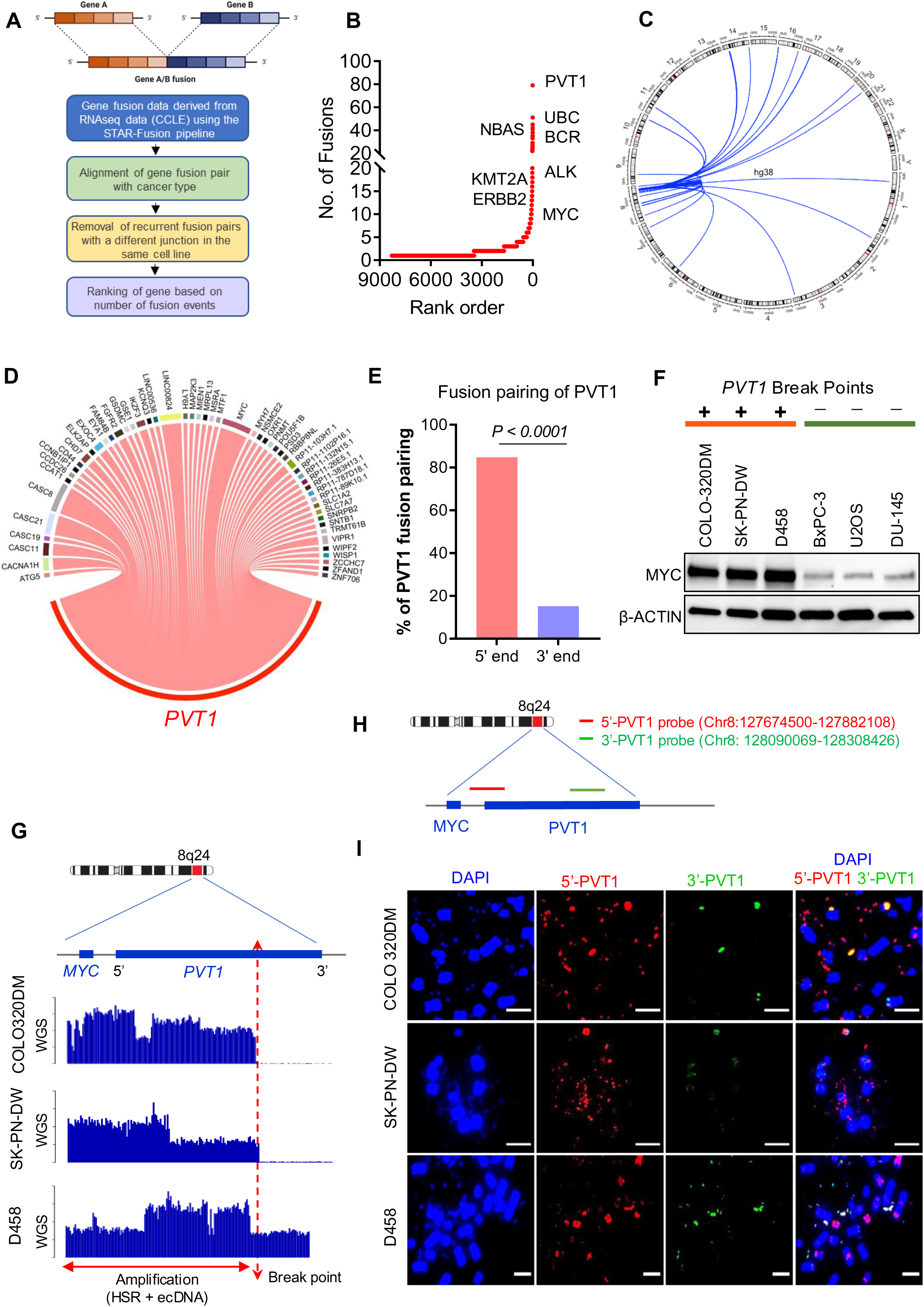
Asymmetric rearrangements of *PVT1* locus in human cancer cell lines: (A) Schematic of gene fusion data analysis using cancer cell line encyclopedia (CCLE) database (B) Genes ranked by unique fusions (x-axis, lowest rank = highest number of unique fusions) plotted against total number of unique fusions in CCLE database (y-axis). The most common participants of recurrent gene fusions are highlighted by their name. (C) A Circos plot representing inter and intrachromosomal *PVT1* fusions. Human chromosomes are radially represented proportional to size, and individual lines represent the *PVT1* locus on 8q24 linking to the location of the corresponding *PVT1* fusion partner. (D) Diverse partners of *PVT1* are shown in the Sankey plot, with the thickness of the connecting ribbon representing the number of times each fusion is observed in CCLE. (E) Percentage of total fusion events showing the proportion of *PVT1* fusions with *PVT1* as the 5’ partner or 3’ partner (P value by Fisher’s exact test). (F) Western blot analysis of c-MYC expression in human cancer cell lines harboring genomic breakpoints in the *PVT1* locus (COLO320DM, SK-PN-DW and D458) and cancer cell lines without genomic breakpoints in the *PVT1* locus (BxPC-3, U2OS and DU145). (G) Genomic copy number profile of COLO320DM, SK-PN-DW and D458 across the MYC-PVT1 region using whole genome sequencing analysis. (H) Schematic of dual-colored break-apart FISH probes for *PVT1* locus; (I) Representative dual-color fluorescence in situ hybridization (FISH) of COLO320DM, SK-PN-DW and D458 cells during metaphase using DAPI and probes spanning the 5’ part of *PVT1* (Red), 3’ part of *PVT1* (Green) and both (merge), Scale bars 10 µm. All imaging experiments were repeated at least three times, with similar results.

To further investigate this phenomenon, we examined three diverse cancer cell lines that harbored MYC/PVT1 amplification and PVT1 translocations: COLO-320DM (colorectal cancer), SK-PN-DW (primitive neuroectodermal tumor (PNET)), and D458 (medulloblastoma (MB)). We also included MYC/PVT1-neutral cell lines (no amplification and no translocation): U2OS (osteosarcoma), BxPC-3 (pancreatic ductal adenocarcinoma) and DU145 (prostate cancer) as controls. We found that the cell lines with amplification of MYC/PVT1 and PVT1 rearrangements (COLO-320DM, SK-PN-DW, and D458) had substantially elevated MYC protein levels, compared to cell lines that did not have MYC/PVT1 amplification or rearrangements (Figure 1F). Whole-genome sequencing (WGS) data analysis confirmed that these cell lines exhibited co-amplification of MYC and 5’-PVT1, along with a significant reduction in 3’-PVT1 genomic content (Figure 1G). Dual-color fluorescence in-situ hybridization (FISH) using region-specific probes also corroborated the selective enrichment of 5’-PVT1 in these cell lines (Figure 1H). COLO-320DM and SK-PN-DW almost exclusively harbored the 5’-PVT1 genomic amplification in the form of ecDNAs, while D458 showed 5’-PVT1 in both ecDNA and HSR amplifications (Figure 1I). These findings collectively substantiate that genomic rearrangements at the PVT1 locus consistently induce asymmetric enrichment of 5’-PVT1, with concomitant loss of 3’-PVT1, underscoring a complex, but potentially critical mechanism in oncogenesis.

### Firefox (FFX): A peptide generated by circularization of PVT1 Exon 2

The abundance of transcripts from 5’-PVT1 in cancer cells suggested that 5’-PVT1 might play a crucial oncogenic role in these cell lines. Although the primary PVT1 transcript contains nine exons, the Ensembl database suggests that the human PVT1 locus can generate 190 transcript variants (<u>ENSG00000249859</u>), which we broadly classified into four sub-categories (Supplementary Figure 2, Supplementary Table 6): variant subclass 1 containing full length *PVT1*, variant subclass 2 containing exons 1, 2 and 3A, and similar iterations spanning *5’-PVT1*; variant subclass 3 containing exons 4A-9 (and their different sub-iterations), and variant subclass 4 containing exons 6A-9. Subclasses 3 and 4 span the *3*’ region of *PVT1*. We hypothesized that these different sets of transcripts might have distinct roles in tumorigenesis.

We have previously shown that siRNA against exon 2 of PVT1 can result in down-regulation of MYC ^8^. A unique feature of *PVT1* exon 2 is that this exon can generate a circular RNA variant of PVT1 (CircPVT1) ^17 18^ (Figure 2A, Supplementary Figure 3 A-C). CircPVT1 expression has been associated with MYC+ cancers, but its functional role in tumorigenesis has remained unclear ^19 20^. We found that even though PVT1 is considered a non-coding RNA, circularization of exon 2 creates an open reading frame encoding a 104 amino acid long peptide, which we termed Firefox (FFX, Figure 2 A, B, Supplementary Figure 3 D). To confirm the endogenous expression of FFX in human cells, we used CRISPR/Cas9 gene editing in U2OS cells to append an in-frame sequence coding for HiBiT ^21^, a small, 11 amino acid peptide, just before the FFX stop codon (Figure 2C). HiBiT binds with high affinity to LgBiT, and the bound complex has high luciferase activity in the presence of added substrate. We carried out a bioluminescence assay to confirm endogenous expression of FFX in several engineered clones (Figure 2D). Sequencing of the luciferase-positive clones confirmed the in-frame knock-in of the HiBiT sequence before the FFX stop codon (Supplementary Figure 4A). We also carried out immunofluorescence microscopy using a HiBiT antibody, which can be used to detect expression of HiBiT-tagged FFX in these engineered cells (Figure 2E, Supplementary Figure 4B). Furthermore, an analysis of human proteome databases identified peptide sequences unique to the FFX peptide (Supplementary Figure 5 A-C, Supplementary Table 7). Notably, one of the identified sequences in the proteomics database, MHVPSGAQLGRPDLLAR, spans the junction sequence of CircPVT1 (connecting the 3’ and 5’ ends of Exon 2), thus providing strong evidence that FFX is encoded by CircPVT1. These experiments collectively confirm the endogenous expression of FFX in human cells.

**Figure 2:**
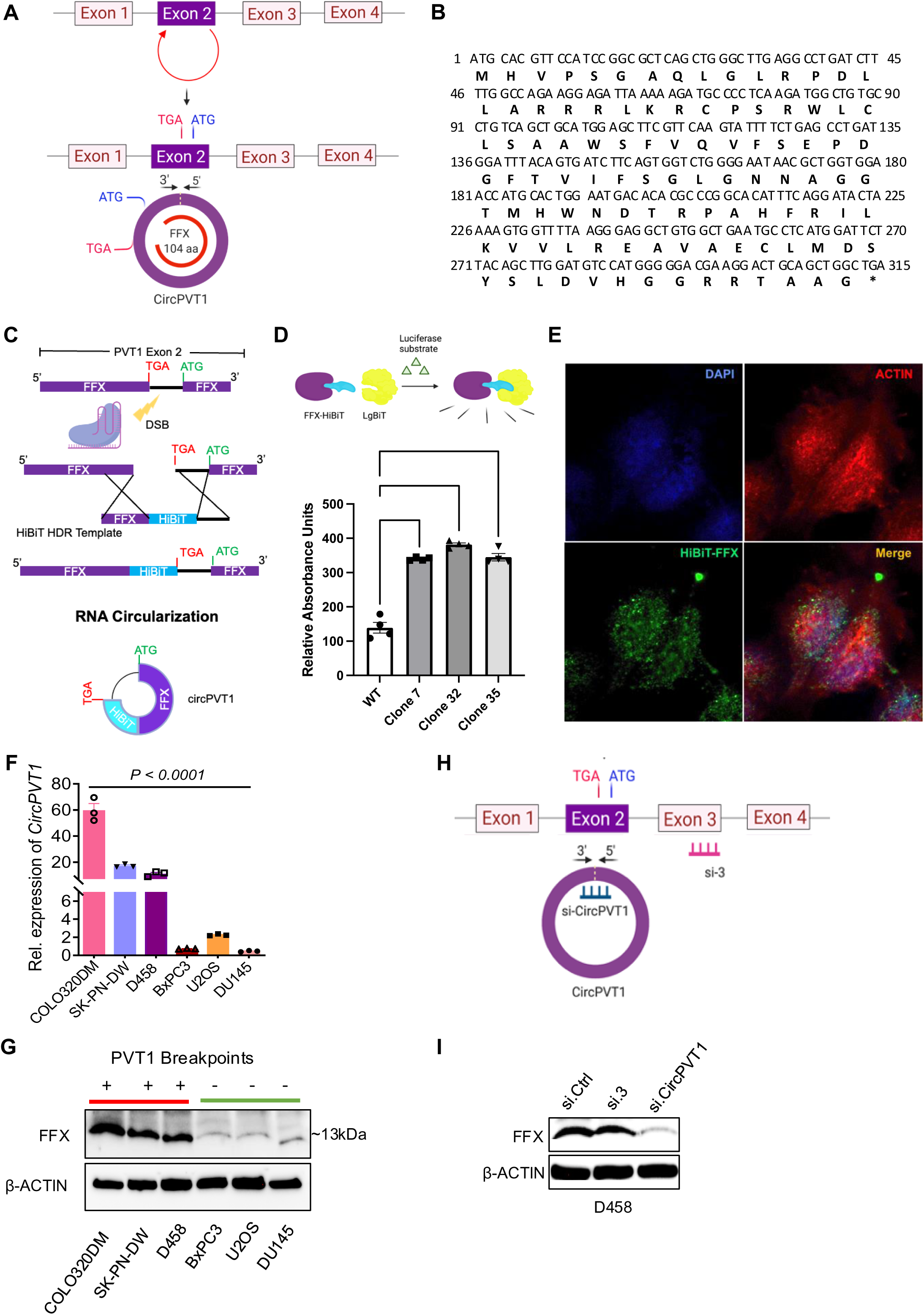
Circularization of *PVT1* exon 2 generates a novel protein Firefox (FFX): (A) Schematic of generation of *CircPVT1* from *PVT1* exon 2 (upper panel), and subsequent re-organization of its open reading frame (ORF), coding a 104 amino acid long peptide (FFX). (B) Coding sequence of FFX. (C) Schematic showing integration of HiBiT sequence in endogenous loci of FFX (immediately upstream of native stop codon to generate C-terminal protein fusion) using CRISPR/Cas9 Knock-in. (D) A HiBiT/LgBiT complementation bioluminescence assay to confirm endogenous expression of FFX in several engineered clones expressing FFX fused with HiBiT peptide. (n=4 P value by on-way ANOVA). (E) Immunofluorescent detection of HiBiT-tagged FFX in CRISPR-edited cell clones using the Anti-HiBiT monoclonal antibody (Green), DAPI (blue) and Actin (red). (F) Quantitative RT-PCR of *CircPVT1* expression in COLO320DM, SK-PN-DW and D458 and *MYC/PVT1* neutral (BxPC-3, U2OS and DU145) cell lines (n=3; P value by one-way ANOVA) (G) Western blot analysis of FFX in COLO320DM, SK-PN-DW and D458 and *MYC/PVT1* neutral (BxPC-3, U2OS and DU145) cell lines using monoclonal anti-FFX antibody. (H) Schematic of siRNAs against *CircPVT1* and linear *PVT1* (I) Western blot analysis of FFX in D458 using anti-FFX antibody in D458 treated with siRNA for control (*si.Ctrl*), linear (si.3) and *CircPVT1* (*si.CircPVT1*).

We used RT-PCR primers specific for CircPVT1 to demonstrate that it was significantly more abundant in cancer cell lines with MYC/PVT1 genomic rearrangements (MYC/PVT1 gain + PVT1 translocation, COLO-320DM, SK-PN-DW, and D458) compared to MYC/PVT1 neutral cell lines (U2OS, BxPC-3, and DU145) (Figure 2F). Additionally, we developed a mouse monoclonal antibody specific for FFX (anti-FFX). Using the anti-FFX antibody, we detected increased expression of FFX in COLO-320DM, SK-PN-DW, and D458 cell lines compared to MYC/PVT1 neutral cell lines (Figure 2G). To test whether FFX is generated from CircPVT1 or from a linear PVT1 transcript, we used siRNA targeting the junction sequence of CircPVT1 to knock down its expression (si.CircPVT1) and one against exon 3 to knock down the expression of linear PVT1 (si.3) in D458 cells (Figure 2H, Supplementary Figure 6). Whereas siRNA-mediated knockdown of linear PVT1 did not affect the expression of FFX, inhibition of CircPVT1 significantly reduced the protein, establishing that CircPVT1 codes for this peptide (Figure 2I). These results not only provide strong evidence that CircPVT1 codes for a previously-unknown peptide, which we named FFX, but also challenges the concept that PVT1 is a non-coding RNA.

### FFX is a novel modulator AKT signaling in MYC+ cancer cells

Next, we aimed to elucidate the functional role of CircPVT1/FFX in cell lines with MYC/PVT1 genomic rearrangement. We used si.CircPVT1 to inhibit FFX expression in COLO-320DM, SK-PN-DW, and D458 cell lines, and observed reduced cellular proliferation in all three cell lines (Figure 3A, Supplementary Figure 7 A, B). Western blot analyses revealed a significant reduction of MYC protein following si.CircPVT1 treatment in all three cell lines, which suggests that FFX knockdown can impact MYC protein levels (Figure 3B, Supplementary Figure 7 C,D). To further investigate the impact of FFX in these cell lines, we conducted RNA-seq analyses following FFX knockdown. We hypothesized that if FFX regulates MYC signaling, all three cell lines treated with si.CircPVT1 would display reduced expression of MYC target genes. In line with our prediction, both global hallmark pathway analysis (Supplementary Figure 8 A-C) and individual gene set enrichment analysis (Figure 3 C,D, Supplementary Figure 9 A-D) confirmed the depletion of MYC targets across all si.CircPVT1-treated cell lines. These findings thereby implicate FFX as a requisite for MYC activity in these cells.

**Figure 3:**
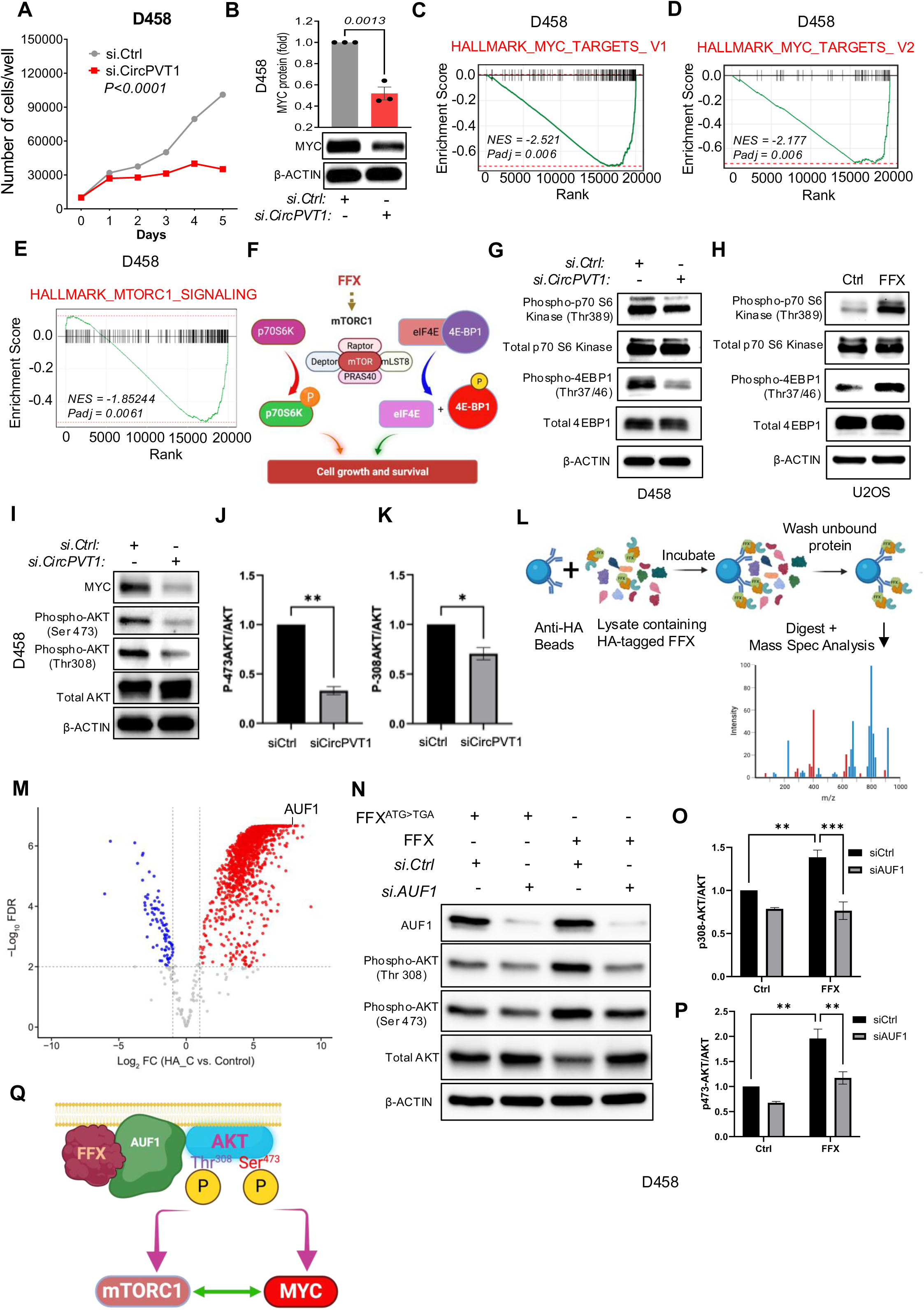
FFX activates AKT signaling in MYC driven cancers: (A) Cell proliferation assay of D458 after transfection with *si.CircPVT1* or si.Ctrl (n=3, P value by two-way ANOVA) (B) Western blot analysis of MYC expression in D458 transfected with si.Ctrl or *si.CircPVT1*. β ACTIN was used as loading control (n=3, P value by unpaired t test). (C, D) Gene set enrichment analysis scores (GSEA) analysis for MYC targets genes in D458 treated with *si.CircPVT1*. *si.Ctrl* was used as control. n=3, for each treatment condition. (E) GSEA of genes involved in MTORC1 signaling in D458 treated with si.CircPVT1. si.Ctrl was used as control. n=3, for each treatment condition. (F) Schematic of the predicted role of FFX in activating the mTORC1 complex (made up of mTOR, Raptor, mLST8, and PRAS40) and the phosphorylation of two key effectors, p70S6 Kinase (p70S6K) and eIF4E Binding Protein 1 (4E-BP1), associated with increased cell growth and survival. (G,H) Western blot analysis of phospho-p70 S6, total p70S6, phospho-4EBP1, total 4EBP1 in (G) D458 transduced with *si.CircPVT1* and *si.Ctrl,* and (H) U2OS overexpressing FFX. A mutated version of FFX (FFX^ATG>TGA^) was served as control. β ACTIN was used as loading control. (I-K) Western blot and quantitative analysis of MYC, phospho-AKT (Ser 473 and Thr308), total AKT expression in D458 cells transduced with *si.CircPVT1* and *si.Ctrl*. β ACTIN was used as a control (p-values obtained by unpaired t-test, n=3, for each treatment condition). (L) Schematic representation of proteomic analysis to identify FFX interacting proteins in U2OS transduced with HA-tagged FFX cDNA. (M) Volcano plot illustrating the significant differential abundance of proteins following co-immunoprecipitation (Co-IP) of protein lysates from U2OS cells transduced with cDNA expressing HA-tagged FFX (HA_C) versus untagged FFX (used as control), using an HA-specific antibody. Proteins with increased abundance post-Co-IP are denoted by red points, while those with decreased abundance are represented by blue points. Proteins were organized by log_2_ fold change (x-axis) and -log10 false discovery rate (FDR) (y-axis). The cutoff lines (-log_10_FDR value of 2 and log2 fold change ± 1) are shown in grey. AUF1 protein is marked exhibiting log_2_fold change of 7.77 and -log_10_FDR of 6.67 (p-value of 2.16e-7). (N) Western blot analysis of AUF1, phospho-AKT (Thr308 and Ser 473), total AKT expression in D458 cells transduced with FFX^ATG>TGA^ (Control) and FFX, and treated with *si.AUF1* and *si.Ctrl*. β ACTIN was used as a control. (O,P) Quantitative analysis of phospho-AKT (Thr308 and Ser 473) expression in D458 cells transduced with FFX^ATG>TGA^ (Control) and FFX, and treated with *si.AUF1* and *si.Ctrl* (p-values obtained by unpaired t-test, n=3, for each treatment condition). (Q) Proposed model for FFX-mediated activation of AKT signaling and the synergistic interaction between mTORC1 and MYC in MYC-driven cancers.

To further validate the functional effects of FFX on MYC expression, we introduced FFX exogenously into MYC/PVT1-neutral U2OS cells. We observed a significant increase in MYC expression following ectopic expression of FFX relative to the control condition (Supplementary Figure 10 A). To distinguish whether it is the CircPVT1 RNA or the encoded peptide FFX that enhances MYC expression, we altered the mRNA sequence of FFX while preserving its amino acid sequence (FFX (CO)). We also generated a FFX mutant, where the start codon ATG was mutated to TGA (FFX^ATG>TGA^), rendering the ORF incapable of coding for the FFX protein, but preserving the remaining mRNA sequence. We transduced U2OS cell line with these DNA constructs and assessed the effects on MYC expression (Supplementary Figure 10 B). The FFX (CO) construct, despite the altered mRNA sequence, significantly augmented MYC protein expression, whereas FFX^ATG>TGA^ did not (Supplementary Figure 10 C,D). This firmly suggests that the FFX protein, not the RNA, has the capacity to enhance MYC levels in these cancer cells.

RNA-seq analysis also indicated a decrease in mammalian target of rapamycin complex 1 (mTORC1) signaling following FFX inhibition in all three cell lines (COLO-320DM, SK-PN-DW, and D458) as demonstrated by global hallmark pathway analysis (Supplementary Figure 8 A-C) and individual gene set enrichment analysis (Figure 3 E, Supplementary Figure 11 A, B). mTORC1 is a critical component of the mTOR signaling pathway, a central regulator of various cellular processes, including proliferation, metabolism, protein synthesis, and cell survival, and is frequently activated in human cancers ^22 23^. To ascertain whether FFX promotes mTORC1 signaling in cancer, we conducted Western blot analyses on two key mTORC1 substrates, p70S6K and 4E-BP1 (Figure 3F), in cell lines where FFX expression was either inhibited or enhanced. FFX inhibition in D458 resulted in a notable decrease in the phosphorylation of p70S6 Kinase (Thr389) and 4EBP1 (Thr37/46) (Figure 3G), whereas FFX overexpression in U2OS led to increased phosphorylation of p70S6K and 4E-BP1 (Figure 3H). These results provide strong evidence that FFX is integral to mTORC1 signaling in human cancer. Thus, we conclude that FFX potentiates MYC and augments mTOR signaling in human cancer.

A mechanism that commonly regulates both MYC and mTORC1 signaling in cancer is the activation of the PI3K/AKT pathway^24, 25^. We therefore investigated if FFX plays a role in the potentiation of the AKT pathway. AKT activation involves the phosphorylation of Serine 473 (Ser473) and Threonine 308 (Thr308) residues within the kinase ^26 27^. Western blot analysis demonstrated that FFX inhibition can lead to a reduction of phospho-AKT^Ser473^, as well as phospho-AKT^Thr308^ in D458 (Figure 3 I-K) and COLO320DM (Supplementary Figure 12). This provides strong evidence that FFX can regulate mTOR and MYC signaling in cancer cells via activation of the AKT pathway. To understand the mechanistic underpinnings by which FFX activates the AKT phosphorylation, we employed a targeted co-immunoprecipitation assay of FFX followed by high-resolution mass spectrometry analysis. We engineered U2OS cells to express HA-tagged FFX, and used anti-HA beads to pulldown FFX and FFX-interacting proteins. The resultant eluate was subjected to comprehensive mass spectrometry analysis (Figure 3L).

Proteomic analysis revealed a plethora of interacting proteins, with AUF1 (HNRNPD) emerging as the second most abundant protein within the eluate ((p-value of 2.16e-7) (Figure 3M, Supplementary Table 8). Western blot analysis on the immunoprecipitated eluate confirmed the association of AUF1 with FFX (Supplementary Figure 13A). Recent studies have shown that AUF1 can localize to the inner cell membrane where it facilitates the phosphorylation and activation of AKT ^28 29^. To determine AUF1’s role in FFX-mediated AKT phosphorylation, we transfected FFX-overexpressing and control cells with siRNA targeting AUF1. Western blots of sample lysate revealed that while cells transduced with FFX exhibited a marked increase in phosphorylation of AKT^Thr308^ as well as AKT^Ser473^, RNAi mediated knockdown of AUF1 in FFX+ cells abrogated the observed increase in phosphor-AKT^Thr308^ and AKT^Ser473^, returning phosphorylation levels of AKT to those seen in the control cells (Figure 3N-P, Supplementary Figure 13 B-D). These findings show that FFX is a key mediator in the activation of AKT by AUF1 in MYC+ cancers. We propose that FFX orchestrates AUF1’s inner membrane recruitment, facilitating activation of AKT signaling, and thereby mTORC1 and MYC in MYC+ cancer (Figure 3Q).

### Honeybadger, a tumor suppressive micro-peptide encoded by PVT1_ts_ regulates MYC expression

We next investigated the functional consequences of asymmetric loss of genomic material from the distal end of PVT1 (*3’-PVT1*) following rearrangement of the PVT1 locus. We hypothesized that loss of the genomic material, and hence the transcript variant(s) at the 3’ end of PVT1, might be advantageous for cancer cells. Since PVT1 rearrangements are heterogeneous, we sought cell lines with MYC/PVT1 gain + PVT1 translocation, but where only a small portion of 3’-PVT1 was lost. We found the breast cancer cell line SKBR-3 to have such a rearrangement, with a loss of exons 7, 8, and 9 of PVT1 (Figure 4A). We designed siRNAs (si.PVT1_ts_) that would specifically target the variants containing exons 7-9 and we characterized them as *PVT1_ts_*. Indeed, si.*PVT1_ts_* treated *MYC/PVT1* neutral cell lines (U2OS, BxPC-3, and DU-145) proliferate faster (Supplementary Figure 14), and Western blot analysis of si.PVT1_ts_ treated cell lines showed an increase in MYC protein levels. (Figure 4B).

**Figure 4:**
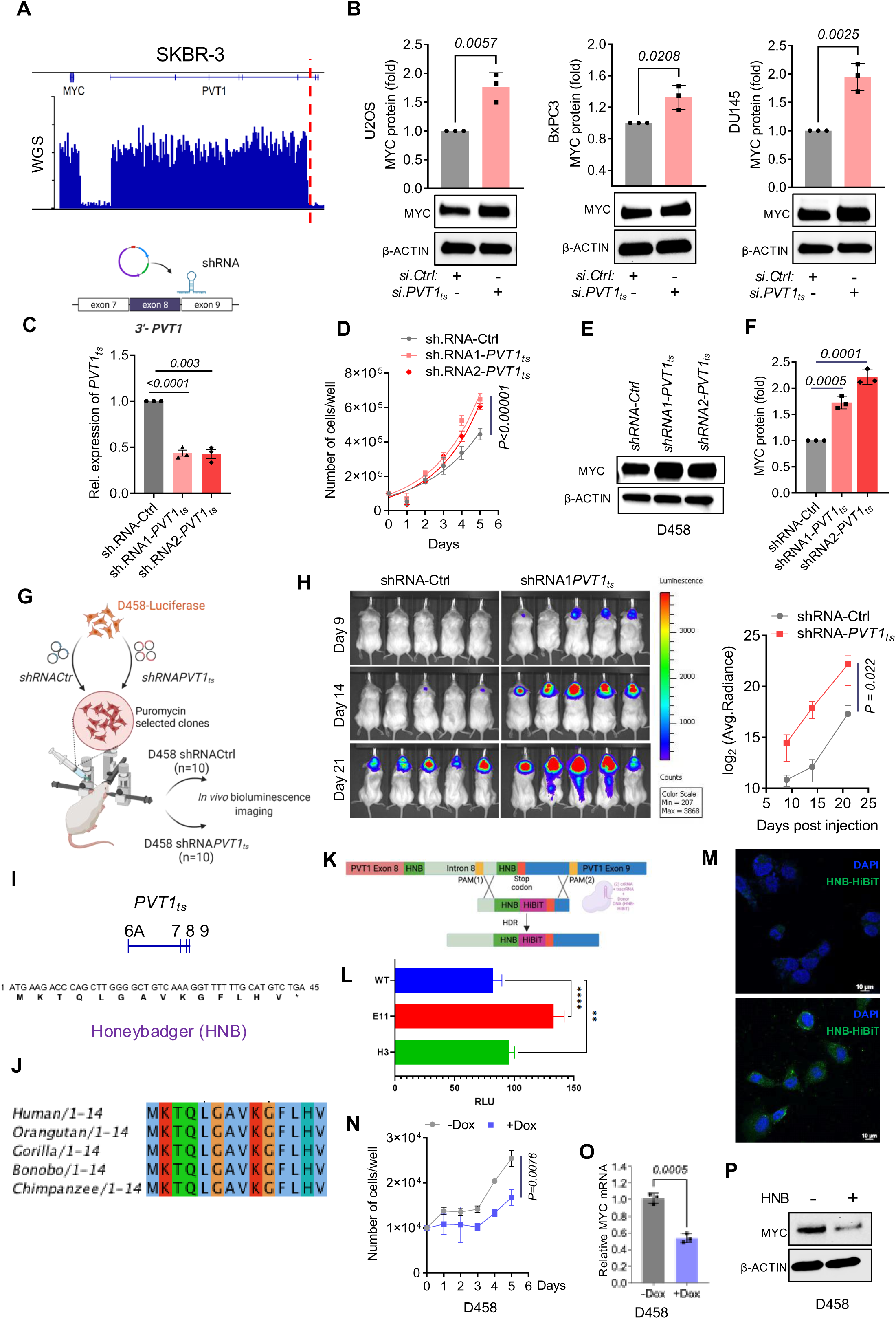
The 3’-PVT1 encodes Honeybadger (HNB), a micropeptide that regulates MYC: (A) WGS coverage of SKBR-3 cells across the MYC-PVT1 region. The translocation at PVT1 is shown by dotted red line (B) Western blot analysis and quantification of MYC expression of si.Ctrl and si.PVT1ts transfected U2Os, BxPC-3 and DU-145 (p-values obtained by unpaired t-test, n=3, for each treatment condition). (C) q-RT-PCR of PVT1ts transcript from D458 transduced with sh.Cntrl, shRNA1-PVT1ts, and sh.RNA2-PVT1ts. (D) Proliferation assay of D458 cells transduced with sh.Cntrl, sh.RNA1-PVT1ts and sh.RNA2-PVT1ts (P values obtained by two-way ANOVA) (E, F) Western blot analysis and quantitation of MYC protein in D458 cells stably transduced with sh.Cntrl, sh.RNA1-PVT1_ts_ and sh.RNA2-PVT1_ts_. β ACTIN was used as a control (p-values obtained by unpaired t-test, n=3, for each treatment condition). (G) Schematic of the orthotropic transplantation assay of D458 cells transduced with sh.Cntrl and sh.RNA1-PVT1ts (H) Bioluminescence assay and quantification of orthotopically transplanted mice at day 9, 14, and 21 (post-transplantation) (n=10 for each of sh.RNA-Cntrl and sh.RNA1-PVT1ts, p values obtained by unpaired t-test). (I) Amino acid sequence of Honeybadger, a micropeptide coded by PVT1ts (J) Sequence alignment of Human HNB showing conserved sequence with other non-human primates. (K) Schematic of CRISPR mediated knock-in of HiBiT tag. (L) HNB-HiBiT lytic detection assay (p-values were obtained One-way ANOVA). (M) Immunofluorescence depicting U2OS cells with sham CRISPR/Cas9 treatment (untagged HNB, serving as control, upper panel) and HNB-tagged HiBiT U2OS cells (lower panel) using Alexa-Fluor 488 for HNB-HiBiT (green) and DAPI (blue) stain for the nucleus. Scale bars represent 10 µm (N) Proliferation assay of D458 following induction of HNB (+/- Dox). (O) q-RT-PCR analysis of MYC transcript levels, with or without induction of HNB (-/+ Dox) in D458 (n=3, P values obtained through unpaired t-test). (P) Western blot analysis of MYC protein expressed in D458 transduced with inducible HNB transgene (-/+ Dox). β ACTIN was used as the loading control

We also reasoned that loss of the residual PVT1_ts_ in cells with MYC/PVT1 gain + PVT1 translocation, or cells with MYC/PVT1 gain without PVT1 translocation, would further augment MYC expression and contribute to increased aggressiveness in these cancer cells. To test this hypothesis, we depleted the residual PVT1_ts_ in D458 cells using two independent shRNAs against *PVT1_ts_* (*shRNA1-PVT1_ts_* and *shRNA2-PVT1_ts_,* Figure 4C). Knockdown of *PVT1_t_*_s_ in D458 cells increased proliferation (Figure 4D) and MYC protein levels (Figure 4 E,F). When D458 cells harboring *shPVT1_ts_* were orthotopically injected into the brains of immunocompromised mice, they grew significantly more aggressively than controls (Figure 4G, H). These results provide strong evidence that a variant of *PVT1 (PVT1_ts_)* at the distal end of the *PVT1* gene, which is depleted in cancer cells harboring *PVT1* rearrangements, encodes a tumor suppressor.

A close inspection of the sequence of *PVT1_ts_* revealed a short open reading frame (sORF) in the last two exons of *PVT1_ts_*. This sORF codes for a 14 amino acid micro-peptide that we named Honeybadger (HNB) (Figure 4I, Supplementary Table 9). HNB is highly conserved in human and several non-human primates (Figure 4J). To confirm the endogenous expression of HNB in human cells, we used CRISPR/Cas9 gene editing to knock-in an in-frame sequence coding for HiBiT immediately before the HNB stop codon in U2OS cells (Figure 4K). We carried out a bioluminescence assay to confirm endogenous expression of HNB, as evidenced by the presence of a signal from the HiBiT protein in two independent engineered clones (Figure 4L). Genomic and RNA sequencing of the luciferase-positive cell lines confirmed the in-frame knock-in of the HiBiT sequence before the HNB stop codon (Supplementary Figure 15 A-C). This suggests that the HiBiT sequence was inserted into the HNB gene in the correct order, without disrupting the reading frame of the gene. Additionally, we carried out immunofluorescence microscopy using HiBiT antibody, which showed that the HiBiT antibody specifically labeled the engineered clones, confirming that they expressed HNB. (Figure 4M). We also identified HNB amino acid sequences with high certainty in several human proteome databases (Supplementary Figure 16, Supplementary Table 10). These results provide strong evidence that HNB is endogenously expressed in human cells.

To test the tumor suppressive function of HNB, we cloned the HNB cDNA in a lentiviral vector with a Doxycycline-inducible promoter. We verified the inducible expression of HNB from this vector by quantitative-RT-PCR following doxycycline-mediated induction (Supplementary Figure 17 A). Induction of HNB expression significantly reduced cellular proliferation of COLO-320DM and D458 cells (Supplementary Figure 17B, Figure 4N). Furthermore, expression of HNB significantly reduced MYC expression at the mRNA as well as the protein levels in these cells (Figure 4 O-P, Supplementary Figure 17 C-D). These data collectively suggest that 3’-PVT1 codes for an anti-tumor micropeptide that regulates MYC. Loss of 3’-PVT1 due to translocation at PVT1 results in depletion of HNB, which further contributes to dysregulation of MYC.

### HNB regulates the RAS/MAPK pathway

To understand the mechanism by which HNB imparts its tumor-suppressive function, we carried out transcriptomic analysis of D458 cells following knockdown of HNB using shRNA against *PVT1_ts_*. Global hallmark pathway analysis (Figure 5A) and GSEA (Figure 5B) revealed that the genes down-regulated by KRAS activation were significantly diminished in HNB-depleted D458 cells. This suggested that HNB may regulate MYC expression via KRAS signaling, a critical component of the mitogen-activated protein kinase (RAS/MAPK) signaling pathway (Figure 5C). Indeed, knocking down HNB in D458 cells augmented RAS/MAPK signaling as evidenced by increased phosphorylation of the downstream effectors of the RAS/MAPK pathway, namely dual specificity mitogen-activated protein kinase kinase 1 and 2 (MEK1/2), extracellular signal-regulated kinases 1 and 2 (ERK1/2), and p90 ribosomal S6 kinase (RSK). (Figure 5D). Similarly, knockdown of HNB in a cell line with MYC/PVT1 amplification but without PVT1 translocation (NCI-H1792, Supplementary Figure 18 A,B) showed similar activation of the RAS/MAPK pathway (Supplementary Figure 18 C, D).

**Figure 5:**
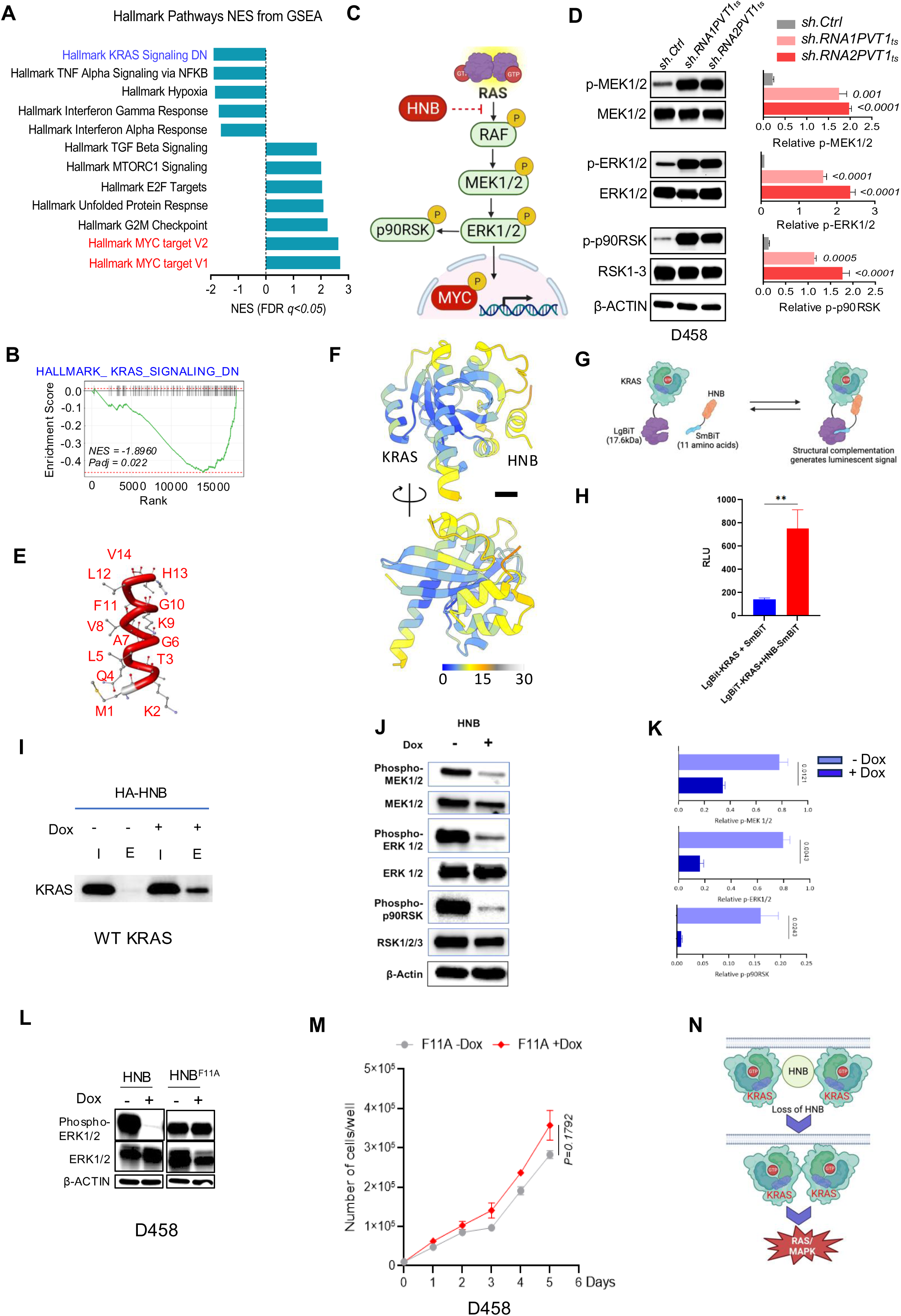
Honeybadger regulates RAS/MAPK pathway: (A) Global hallmark pathway analysis of transcriptome from shRNA1-PVT1ts transduced D458 cell line. sh.Cntrl was used as control (n=3). (B) GSEA analysis for genes downregulated due to KRAS activation in D458 stably transduced with shRNA1-PVT1ts (n=3). (C) Schematic representation of the possible role of HNB in regulating the RAS/MAPK signaling pathway in cancer. (D) Western blot and quantitative analysis of phospho-MEK1/2, total MEK1/2, phospho-ERK1/2, total ERK1/2, phospho-p90RSK, total RSK1-2 expression in D458 transduced with sh.Cntrl, shRNA1-PVT1ts and shRNA2-PVT1ts. β ACTIN was used as a control (p-values obtained by unpaired t-test, n=3, for each treatment condition). (E) Structural model of HNB peptide (Tube stick model). (F) Per-residue predicted aligned error of the AlphaFold prediction between KRAS and HNB. Scale bar: 5 Å. (G) Schematic representation of NanoBiT: protein-protein interaction system with LgBiT tagged KRAS and SmBiT tagged HNB. The interaction of fusion partners leads to the complementation of LgBiT and SmBiT, generating an enzyme with a bright, luminescent signal. (H) Luminescence signal from structural complementation of LgBiT-KRAS and HNB-SmBiT showing an interaction between KRAS and HNB in HEK293T cells. Non-fused HiBiT and LgBiT KRAS were used as control (p-values obtained by unpaired t-test, n=3). (I) Western blot analysis of Co-IP experiments from U2OS transfected with uninduced (-) and doxycycline induced expression of HA tagged HNB (HA-HNB) with KRAS antibody. (J,K) Western blot and quantitative analysis of phospho-MEK1/2, total MEK1/2, phospho-ERK1/2, total ERK1/2, phospho-p90RSK, total RSK 1-3 in D458 transduced with doxycycline-inducible HNB. β ACTIN was used as a control (p-values obtained by unpaired t-test, n=3, for each treatment condition). (L) Western blot analysis of phospho-ERK1/2 and total ERK1/2 in D458 transduced with doxycycline-inducible HNB and HNB^K9Q^. β ACTIN was used as a control. (M) Proliferation assay of D458 following induction of HNB^F11A^ (+/- Dox) (N) Graphic representation of oncogenic RAS/MAPK signaling activation due to the loss of HNB resulting from PVT1 rearrangement in MYC+ cancer.

We then investigated how HNB can regulate the RAS/MAPK pathway. Micro-peptides often exert their biological roles by interacting with and modulating larger proteins^30-32^. Given the profound upregulation of RAS/MAPK signaling following inhibition of HNB, we investigated whether HNB could regulate KRAS by directly interacting with it. We used AlphaFold 3 ^33^ to assess the potential protein-protein interaction between HNB (Figure 5E) and KRAS (PDB ID:4OBE). The algorithm predicted that HNB could interact with KRAS with high probability (Figure 5 F, Supplementary Figure 19). To confirm the intracellular interaction of KRAS and HNB, we employed a NanoBiT complementation reporter assay ^34^ where KRAS was fused to a larger subunit (LgBiT) of a NanoLuc luciferase, and HNB was fused to a smaller subunit (SmBiT) or a HiBiT fragment. A physical interaction between KRAS and HNB would bring LgBiT and SmBiT (or HiBiT) in proximity, generating a luminescent signal (Figure 5G, Supplementary Figure 20 A). As predicted, HNB-SmBiT and HNB-HiBiT fragments exhibited significant increase in bioluminescence with KRAS-LgBiT, as compared to the fragments alone (Figure 5H, Supplementary Figure 20 B). To quantify the interaction thermodynamics, we performed isothermal titration calorimetry (ITC) ^35^. While HNB self-interaction was not detected (Supplementary Figure 21 A,B), titration of HNB into KRAS-containing buffer yielded a clear binding signal with a Kd of 364 ± 2 µM, indicating a direct interaction (Supplementary Figure 21 C-E), Finally, we performed a co-immunoprecipitation experiment using a doxycycline-inducible HA-tagged HNB construct to investigate the interaction between HNB and KRAS. Western blot analysis confirmed successful co-immunoprecipitation of KRAS following doxycycline-induced expression of HA-tagged HNB in U2OS cells (Figure 5I). Collectively, these data support our hypothesis of an intracellular interaction between KRAS and HNB.

We next tested whether HNB could interfere with KRAS dimerization, a pivotal event in the activation of the RAS-MAPK signaling cascade ^36-38^ We used an acceptor photobleaching assay, a specialized variant of Fluorescence Resonance Energy Transfer (FRET) used to study the interaction between two biological molecules ^39^. Typically, in a FRET experiment, a “donor” protein (KRAS-CFP) and an “acceptor” protein (KRAS-YFP) are tagged with distinct fluorescent proteins. When in close proximity, the donor can transfer energy to the acceptor, resulting in fluorescent emission from the latter. In acceptor photobleaching assay, the acceptor fluorophore is photobleached. In the unbleached state, the donor fluorophore’s fluorescence is reduced because it transfers energy to the acceptor. After acceptor photobleaching, the energy transfer stops because the acceptor can no longer absorb energy. As a result, the donor fluorophore’s fluorescence increases (Supplementary Figure 22 A). By comparing the fluorescence of the donor, KRAS-CFP, before and after the photobleaching of the acceptor, either in the presence or absence of HNB, we can infer whether FRET was occurring and, consequently, whether HNB can modulate the interaction between KRAS monomers. We hypothesized that loss of HNB would promote interaction between KRAS monomers (Supplementary Figure 22 B). Consistent with this hypothesis, the acceptor photobleaching assay on D458 cells with depleted HNB displayed a marked increase in CFP (the donor fluorophore) fluorescence post-photobleaching of YFP (the acceptor fluorophore) (Supplementary Figure 22 C,D). This strongly suggests that the absence of HNB facilitates closer proximity between KRAS monomers. Conversely, we also hypothesized that expression of HNB should disrupt the FRET between the KRAS monomers (Supplementary Figure 22 E). Indeed, induction of HNB in shPVT_ts_ treated D458 cells mitigated the augmented CFP fluorescence observed following the photobleaching of YFP, which suggests that HNB can impede the interaction between KRAS monomers (Supplementary Figure 22 F,G). To ascertain whether HNB possesses the ability to modulate the activity of RAS/MAPK pathway effectors, we carried out Western blot analyses of the phosphorylation states of MEK1/2, ERK1/2, and p90RSK in D458 (harboring wild type KRAS) and NCI-H1792 (harboring mutant KRAS^G12C^) cells following the induction of HNB. HNB expression significantly dampened the phosphorylation of the key effectors - MEK1/2, ERK1/2, and p90RSK, (Figure 5 J, K, Supplementary Figure 23) providing strong evidence for the role of HNB as a regulator of RAS/MAPK signaling activity. To further dissect HNB’s capacity to impede RAS-MAPK activation, we postulated that mutating critical residues within HNB would diminish its ability to modulate KRAS activation as well as its tumor suppressive function. Western blot analysis revealed that while ectopic expression of HNB significantly diminished ERK1/2 phosphorylation, the overexpression of an HNB variant carrying a mutation at its F11 residue (HNB^F11A^) failed to inhibit phosphorylation of the same RAS effector (Figure 5L). Additionally, overexpression of HNB^F11A^ failed to suppress the proliferation of D458 (Figure 5M). This observation implies that functional residues within HNB play a crucial role in engaging with KRAS and subsequently regulating RAS MAPK signaling, and mediate HNB’s tumor suppressive function. Collectively, these findings provide compelling evidence that HNB governs KRAS activation, and that the loss of HNB potentiates the activation of the RAS-MAPK pathway (Figure 5N).

### PVT1 rearrangement stabilizes MYC in human cancer

Subsequently, we investigated how genomic alterations at the PVT1 locus can fuel MYC-driven carcinogenesis. Previous studies have implicated the RAS/MAPK signaling pathway in MYC stabilization, as it enhances MYC phosphorylation at Ser62 via phospho-ERK and phospho-RSK signaling pathways^40,41^. We postulated that the loss of HNB due to genomic translocation at PVT1 could result in increased MYC^Ser62^ phosphorylation (Figure 6A). HNB knockdown in D458 and NCI-H1792 cells resulted in elevated phosphorylated MYC^Ser62^ levels (Figure 6B and Supplementary Figure 24 A,B). Cycloheximide chase assays confirmed the enhanced MYC stability upon HNB depletion in both D458 (Figure 6 C,D) and NCI-H1792 cells (Supplementary Figure 24 C,D). RNA-Seq analysis followed by GSEA further corroborated the elevation of MYC targets in HNB-deficient D458 cells (Figure 6 E,F), indicating enhanced MYC activity. Hence, we propose that PVT1 translocations may bolster MYC stability and function via enhanced MYC^Ser62^ phosphorylation, facilitated by HNB loss and subsequent activation of the RAS/MAPK pathway.

**Figure 6.**
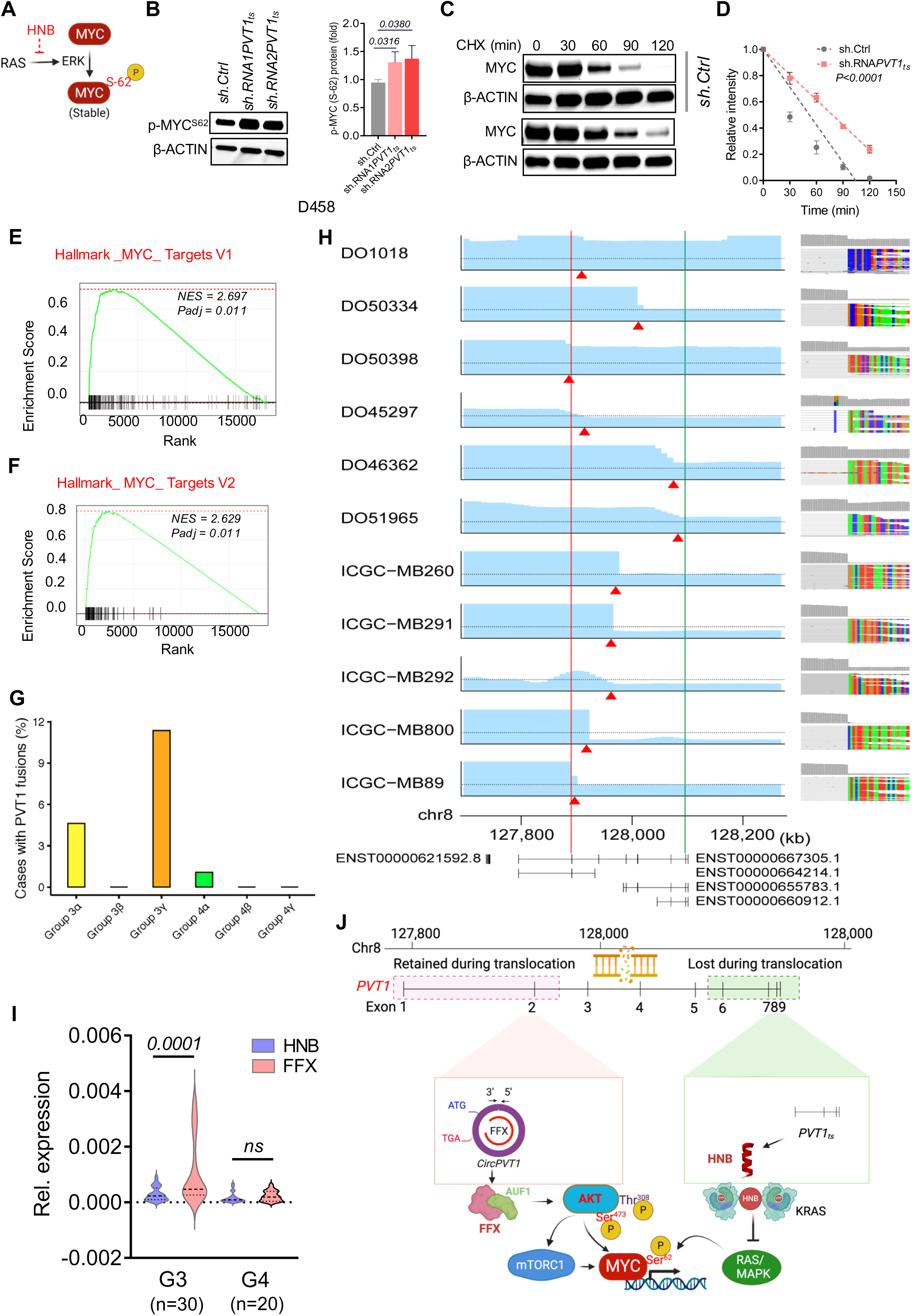
Rearrangements at *PVT1* stabilize MYC in human tumors: (A) Schematic representation of the role of RAS/MAPK signaling in increased phospho-MYCSer62 (p-MYC^Ser62^) (B) Western blot and quantitative analysis of p-MYCSer62 in D458 stably transduced with sh.Cntrl, shRNA1-PVT1ts and shRNA2-PVT1ts. β ACTIN was used as loading control (p-values obtained by ANOVA). (C, D) Cycloheximide chase assay and quantification of c-MYC in D458 cells transduced with sh.Cntrl (as control) and shRNA1PVT1_ts_. β-ACTIN was used as a control (p-value obtained by simple linear regression, n=3, for each treatment condition). (E, F) GSEA analysis of MYC target genes upregulated due to activation of c-MYC in D458 stably transduced with shRNA1-PVT1_ts_ (n=3). (G) PVT1 fusion enrichment analysis from RNASeq data from the MB tumors (N= 545). (H) Pan-cancer analysis of unbalanced PVT1 rearrangement with a genomic breakpoint between FFX and HNB. Bar charts on the left column show copy number changes. The vertical axis indicates the segmented copy number ratio, and the dotted line indicates the number of two. The red triangles correspond to the breakpoints. The chromosomal coordinates from the human genome reference (GRCh38) are shown at the bottom. The red and green lines represent the position of the 3′ end of FFX and the 5′ end of HNB, respectively. Representative isoforms of MYC and PVT1 are also shown from GENCODE v.38. The right column shows the screenshots of the IGV at the breakpoint of each case. (I) q-RT-PCR of FFX (red) and HNB (Blue) transcripts from RNA samples derived from G3 (n=30) and G4 (n=20) MB patients (P value determined by unpaired t-test). (J) Proposed model for how AKT and RAS/MAPK signaling pathways converge, enhancing MYC activation due to rearrangements at the PVT1 locus in MYC-driven cancers.

To assess that, we probed the prevalence of PVT1 fusion events across distinct Medulloblastoma (MB) subgroups. Notably, Group 3 (G3) MB is recognized as MYC-driven and particularly lethal ^42-44^. Our analysis of RNA-Seq data from diverse MB tumor subtypes revealed PVT1 fusions to be most prevalent in G3 gamma, a subtype featuring MYC-PVT1 amplification and PVT1 translocation, as well as in G3 alpha (Figure 6G). This aligns with previous findings of elevated p-Ser^62^ MYC and enhanced MYC activity in G3 gamma and G3 alpha tumors relative to other MB subgroups ^45^.

To examine whether FFX enrichment and HNB loss occur in human tumors, we scrutinized the MYC-PVT1 region in whole genome sequencing (WGS) data from primary human tumors in the ICGC-PCAWG ^46^ and Medulloblastoma ^47^ datasets. We identified several tumors exhibiting PVT1 genomic amplification and rearrangement, characterized by an enrichment of the genomic region encoding FFX and a concomitant loss of the region encoding HNB (Figure 6H). Additionally, we conducted RT-qPCR analysis of FFX and HNB transcripts in RNA samples from 50 human MB tumors. While we observed no difference in expression of FFX and HNB in Group 4 (G4) MB samples, significant enrichment of FFX expression over HNB expression was observed in G3 MB samples (Figure 6I). In conclusion, our data collectively implicate PVT1 as a crucial hub regulating MYC potency by modulating both AKT and MAPK activity in human cancer (Figure 6J). Genomic amplification of MYC/PVT1 locus can facilitate FFX enrichment, which can augment AKT activity, while a genomic translocation at PVT1 can induce loss of HNB, thereby stimulating MAPK signaling in human tumors. We therefore propose that rearrangement at PVT1 is a sophisticated mechanism exploited by cancer cells to sculpt the oncogenic landscape of MYC-driven malignancies.

## DISCUSSION

MYC-driven tumors are a formidable challenge in oncology, characterized by poor prognosis and adverse patient outcomes. The simplistic view of MYC dysregulation does not fully encapsulate the complexity underlying these clinical presentations. Previously, we have proposed a compelling correlation between MYC and PVT1 in 8q24+ tumors ^8^. Additional evidence of PVT1 breakpoints leading to PVT1 promoter-MYC fusions on extrachromosomal circular DNA (ecDNA), fostering unregulated enhancer input to drive MYC transcription have also strengthened the MYC-PVT1 co-operation in MYC+ cancers ^48 49 50 51^. In this work, we reveal that translocation events at the PVT1 locus may act in synergy, orchestrating a convergence of MYC, AKT, mTOR, and MAPK signaling pathways. This ultimately enhances MYC stability and enriches its targets within these malignancies.

Historically categorized as ‘non-coding’ RNA, our findings illuminates the dual functionalities of PVT1 as both oncogenic and tumor-suppressive via newly identified peptides, FFX and HNB, respectively. The discovery aligns with emerging literature, which attributes coding potential to lncRNAs, including micropeptides ^52^, and their transformative role in muscle development, metabolism, and cell signaling. ^53-55^. Similarly, once perceived as non-coding entities, circular RNAs have been found capable of translation into functional peptides. ^56 57^. This adds to the expanding view of lncRNAs in cellular transformation.

We disclose a novel and crucial mechanistic association between FFX and activation of AKT kinase, a central modulator of cell proliferation, growth, and survival, mediated by FFX-AUF1 interaction. Upon activation, AKT can reprogram cellular metabolism, inciting mTORC1 – a pivotal regulator of protein synthesis – through specific phosphorylation of translation initiation factor 4E (eIF4E) binding protein 1 (4EBP1) and ribosomal protein p70S6 kinase (p70S6K1/2) ^58 59 60^. Activated AKT also controls GSK3β, subsequently augmenting MYC activity in cancer cells. Our data substantiate FFX’s essential role in reinforcing the AKT/mTORC1/MYC axis, aligning with prior insights linking MYC and mTOR-dependent regulation of protein synthesis in cancer ^61 62^. While the detailed mechanism by which FFX sensitizes AKT/mTORC1/MYC axis needs to be investigated, the targeting of FFX emerges as a promising therapeutic strategy for MYC-dependent cancers, offering a potential avenue for combating the otherwise elusive direct inhibition of MYC.

Our study also uncovers the implications of the loss of HNB encoded by 3’-PVT1 in the activation of RAS/MAPK signaling. Building on the foundational work that established MYC and RAS as co-operating oncogenes ^63^, we propose that the loss of 3’-PVT1 facilitates an activated RAS/MAPK pathway, prolonging the MYC protein’s half-life and intensifying its activity. Our preliminary insights into HNB-KRAS interaction lay the groundwork for further exploration, potentially leading to innovative therapeutic interventions in RAS and MYC-driven malignancies.

In summary, we identify PVT1 as a complex molecular hub that orchestrates the interplay between MYC, mTOR, AKT, and RAS signaling in cancer. This study not only elucidates intricate regulatory mechanisms that appear to be selectively utilized during tumorigenesis, but also identifies potential therapeutic avenues that could be leveraged for patient benefit.

## MATERIALS AND METHODS

### Cell Lines and Culture Conditions

Cancer cell lines: COLO-320DM, SK-PN-DW, U2OS, BxPC-3, DU145, SKBR-3 and, NCI-H1792 were obtained from American Type Culture Collection (Rockville, MD) and grown in their required growth medium per the American Type Culture Collection description. D458 and MB002 were obtained from Dr. Michael Taylor’s lab at SickKids hospital, Toronto. Cell line identity for the master stocks were verified by STR genotyping (IDEXX BioResearch)

### Western blot Analysis

For whole-cell lysate, the cell pellet was resuspended in RIPA buffer (Cell Signaling Technologies, Danvers, MA) supplemented with a protease inhibitor cocktail (Cell Signaling Technologies) and incubated on ice for 20 minutes. The lysate was then centrifuged at 14,000 revolutions/minute for 10 minutes at 4°C, and the supernatant was harvested. Protein concentration was quantified by BCA Protein Assay Kit (Thermo Fisher Scientific) as per kit protocol. Next, 20 μg of whole-cell lysate was electrophoresed on a 4%–20% gradient polyacrylamide gel with SDS (Bio-Rad, Hercules, CA) and electroblotted onto polyvinylidene difluoride membranes (Bio-Rad). Membranes were blocked in Tris-buffered saline with 5% nonfat milk and 0.1% Tween and probed with antibodies. Bound proteins were detected with horseradish-peroxidase–conjugated secondary antibodies (Cell Signaling Technologies) and SuperSignal West Femto (Thermo Fisher Scientific). Antibodies used were anti-c-MYC (Cell Signaling Technology, 5605, anti-Phospho-c-MYC (Ser62) (Cell Signaling Technology, 13748), anti-β ACTIN (Cell Signaling Technology, 8457), anti-Phospho-p44/42 (Erk1/2) (Thr202/Tyr204) (Cell Signaling Technology, 4370), anti-p44/42 (Erk1/2) (Cell Signaling Technology, 4695), anti-Phospho-MEK1/2 (Ser217/221) (Cell Signaling Technology, 9154), anti-MEK1/2 (Cell Signaling Technology, 9122), anti-Phospho-p90RSK (Ser380) (Cell Signaling Technology, 11989), anti-RSK1/RSK2/RSK3 (Cell Signaling Technology, 9355), Anti-Rabbit IgG, HRP-linked Antibody (Cell Signaling Technology, 7074). Anti-FFX (Firefox) monoclonal antibodies were custom produced by Proteintech using FFX residues 13-104 as epitope.

### Real-Time Polymerase Chain Reaction Analysis

Total RNA was extracted with the TRIzol (Roche, Indianapolis, IN) as described by the manufacturer and complementary DNA synthesis, 1 μg of total RNA was reverse transcribed by using the GoScriptTM Reverse Transcription Mix, Random Primer Protocol (Promega, Madison, WI). Real-time PCR was run in triplicate using PowerUpTM SYBRTM Green Master Mix (Thermo Fisher Scientific), following the manufacturer’s instructions. Real-time PCR amplification monitoring was performed using the CFX96 Touch Real-Time PCR Detection System (Bio-Rad). Data were expressed as relative mRNA levels normalized to the GAPDH, served as endogenous normalization control expression levels in each sample, and are represented as mean ± standard error of the mean between 2 independent experiments unless otherwise indicated in the figure legend. The primer sequences are listed in the Supplementary Table.

### Lentiviral Production and Infection

Lentivirus was produced in HEK293T cells by transfecting plasmids and packaging plasmids (pMD2.G and psPAX2) (Addgene) by using Fugene HD Transfection Reagent (Promega). Media was replaced with R10 media 6–8 hours after transfection, and lentivirus-containing supernatant was subsequently collected every 12 hours for 48 hours before filtration through a 0.45-mm filter. The lentivirus was then concentrated using the Lenti-X Concentrator (Clontech). For infection of cells, the cells were plated at 60% confluency in each well of the 6 well plates. The cells were then infected with 25 µl of concentrated lentivirus and 10 µg/mL of Polybrene. The media was changed 24 hours after transduction. 48 hours after transduction, the cells were selected using a 1 µg/mL concentration of Puromycin.

### Cycloheximide Assay

Cycloheximide (50 μg/mL) was added to the cells 24 hours after seeding. The cells were lysed at indicated time points, and the level of MYC was determined by Western blotting using an anti-MYC antibody. The signal intensity of MYC protein was normalized to β ACTIN. For quantification, the percent change in signal intensity at time 0 was set as 1 hour.

### Doxycycline Inducible Overexpression System

The HNB ORF and eGFP (control) were cloned in an inducible lentivirus expression, TRE-gateway plasmid (pLIX_403, Addgene, #41395). Lentivirus was produced in HEK293T cells using the protocol described above. D458, NCI-H1792 and COLO 320DM cells were transfected with inducible HNB lentivirus and selected with 1 µg/mL concentration of Puromycin. Selected cells were seeded at a concentration of 4 x 10^5^ cells per well. After 24 hours of seeding, the cells were treated with 1 µg/mL Doxycycline for 48 hours. The cells were then lysed using RIPA buffer and 1X protease inhibitor cocktail. The protein lysates were collected and quantified using the previously described protocol. 20 μg of whole-cell lysate was electrophoresed on a 4%–20% gradient polyacrylamide gel with SDS (Bio-Rad, Hercules, CA) and electroblotted onto polyvinylidene difluoride membranes (Bio-Rad). Membranes were blocked in Tris-buffered saline with 5% nonfat milk and 0.1% Tween and probed with antibodies. Bound proteins were detected with horseradish-peroxidase–conjugated secondary antibodies (Cell Signaling Technologies) and SuperSignal West Femto (Thermo Fisher Scientific). Antibodies used were anti-c-MYC (Cell Signaling Technology, 5605, anti-Phospho-c-MYC (Ser62) (Cell Signaling Technology, 13748), anti-p70S6K (Cell signaling Technology, 9202), anti-Phospho-p70S6K (Thr389) (Cell signaling Technology, 92025), anti-4E-BP1 (Cell signaling Technology, 9644), anti-Phospho-4E-BP1 (Thr37/46) (Cell Signaling Technology 2855), anti-AKT (Cell signaling Technology 9272), Anti-Phospho-AKT (Ser473) (Cell Signaling Technology 9271) anti-Phospho-AKT (Thr308) (Cell Signaling Technology 9275) anti-β ACTIN (Cell Signaling Technology, 8457), anti-Phospho-p44/42 (Erk1/2) (Thr202/Tyr204) (Cell Signaling Technology, 4370), anti-p44/42 (Erk1/2) (Cell Signaling Technology, 4695), anti-Phospho-MEK1/2 (Ser217/221) (Cell Signaling Technology, 9154), anti-MEK1/2 (Cell Signaling Technology, 9122), anti-Phospho-p90RSK (Ser380) (Cell Signaling Technology, 11989), anti-RSK1/RSK2/RSK3 (Cell Signaling Technology, 9355), Anti-Rabbit IgG, HRP-linked Antibody (Cell Signaling Technology, 7074).

### Gene fusion data from the Cancer Cell Line Encyclopedia (CCLE)

Gene fusion data from CCLE previously published 65 was obtained and processed to determine the number of times each fusion pair appears from RNAseq fusion calling pipelines in CCLE. Fusions were called using three different algorithms: deFuse, TopHat-Fusion, and STAR-Fusion 66 67 68 and stringent filtering criteria were applied to remove false positives and fusion pairs found in normal tissue. Using this data, we removed duplicates (i.e.: same fusion pairs in the same cell line with a different junction) and determined the total number of times each gene is involved in fusion pairs. We then sorted this data by cancer/tissue type and fusion type (i.e. whether the gene is found at the 5’ or 3’ end of the fusion).

### Metaphase chromosome spread

Metaphase spreads were prepared as described before 49 52. Cells in metaphase were prepared by KaryoMAX (Gibco) treatment at 0.1 μg ml−1 for 3 h. Single-cell suspension was then collected and washed by PBS, and treated with 75 mM KCl for 15–30 min. Samples were then fixed by 3:1 methanol:glacial acetic acid, v/v and washed an additional three times with the fixative. Finally, the cell pellet resuspended in the fixative was dropped onto a humidified slide.

### Metaphase DNA FISH

The dual probes specific to 5’-PVT1 half (hg38 /human; chr8:127674500-127882108) was labeled with Cyanine 5 dye (650nm) and 3’-PVT1 half (hg38/human; chr8:128016680-128234071) was labelled with Cyanine 3 dye (550nm). Slides containing fixed cells in metaphase were briefly equilibrated by 2× SSC, followed by dehydration in 70%, 85% and 100% ethanol for 2 min each. FISH probes in hybridization buffer (Empire Genomics) were added onto the slide, and the sample was covered by a coverslip, denatured at 75 °C for 1 min on a hotplate and hybridized at 37 °C overnight. The coverslip was then removed, and the sample was washed once by 0.4× SSC with 0.3% IGEPAL, and twice by 2× SSC with 0.1% IGEPAL, for 2 min each. DNA was stained with DAPI and washed with 2× SSC. Finally, the sample was mounted by mounting medium (Molecular Probes) before imaging.

### Analysis of PVT1 rearrangement by whole-genome sequencing (WGS) data

The WGS cohort includes two datasets, ICGC-PCAWG and medulloblastoma. The medulloblastoma dataset consists of 341 cases from the ICGC and Toronto cohorts used in the previous study. We removed 142 duplicated cases with medulloblastoma from the ICGC-PCAWG dataset. For the ICGC-PCAWG dataset, we investigated the GISTIC copy number result provided by the PCAWG study (https://dcc.icgc.org/releases/PCAWG/consensus_cnv/GISTIC_analysis) to investigate copy number gain of the MYC and/or PVT1 loci. For the cases without GISTIC, we analyzed structural variants on the MYC and/or PVT1 loci. The cases that have copy number gain or breakpoints on the MYC and/or PVT1 loci were analyzed for the change of read coverage between FFX and HNB using IGV.

For the medulloblastoma cohort, we aligned whole-genome sequencing reads to 1000 Genomes Project GRCh38DH human genome reference sequence that includes decoy sequences and alternate versions of the HLA locus (ftp://ftp.1000genomes.ebi.ac.uk/vol1/ftp/technical/reference/GRCh38_reference_genome/GRCh38_full_analysis_set_plus_decoy_hla.fa), using Burrows–Wheeler aligner (BWA) – MEM version 0.7.17 with the default setting. Copy-number alterations were analyzed using Control-FREEC v.11.6. with the following parameters: breakPointType = 2, ploidy = “2,3,4”, step = 10000, window = 50000, and we collected the tumors with focal copy number gain (absolute copy number ≥ 3) of the MYC and/or PVT1 locus from the results. We visually reviewed the change in depth between FFX and HNB. The ICGC cases suspected with a breakpoint between FFX and HNB were remapped using BWA and analyzed for copy number changes with the same methods described above. Structural variant (SV) calling was performed using GenomonSV (https://github.com/ Genomon-Project) with the following parameters: min_overhang_size = 50, sv_min_tumor_allele_freq = 0.02. Detected SVs between FFX and HNB were visually reviewed using the IGV.

### Sequence alignment of HNB in Human and non-human primates

BLAST analysis identified the sORF coding for HNB were found in the following non-human primates: Sumatran Orangutan (XM_024250917.1) Gorilla (XR_004071200.1), Bonobo (XR_004673073.1), Chimpanzee (XR_001720429.2). Clustal W was used to carry out the multiple alignment of the translated sequence of the sORF.

### FRET

pcDNA3.1 CFP-KRAS (#112717) and pcDNA3.1 YFP-KRAS (#112718) were purchased from Addgene. The plasmids were co-transfected into D458 cells in a 35 mm cell culture dish with glass bottom. The cells were imaged live 48 hours after transfection on the Nikon Super Resolution Confocal Microscope. CFP was excited using 405-445nm laser and YFP was excited using the 514 nm laser. YFP was photobleached at 80% power for 1 minute and the CFP and YFP were imaged after photobleaching using the respective laser and filters. The images were analyzed in Fiji, ImageJ.

### Isothermal Titration Calorimetry

Isothermal titration calorimetry (ITC) measures the change in temperature when two or more biomolecules bind to each other in a solution. The assay provides the binding affinity of ligands to proteins and the entropy or enthalpy of the reaction. Recombinant KRAS protein (#ab156968, Abcam) was dialyzed into the ITC assay buffer (20mM Tris pH 8.0, 100mM NaCl, 5mM Beta-mercaptoethanol) and concentrated to about 100 μM. During ITC titration twenty 6 μl injections (with intervals of 150 seconds) of 500μM HNB were made into the ITC cell containing 93 μM KRAS. Titration of 500μM HNB into the ITC assay buffer under the same conditions was used as the baseline control. ITC experiments were performed using Affinity ITC instrument (TA Instruments) at 25°C. The data was fitted and analyzed using NanoAnalyze software provided by TA Instruments.

### NanoBiT Protein: Protein Interaction System

LgBiT-KRAS, HNB-SmBiT, and SmBiT (Control) were cloned into pcDNA3.1 plasmid vectors. HEK293T cells were plated in opaque clear bottom 96-well plates at a confluency of 10,000 cells per well using complete DMEM media (10% FBS and 1% Pen/Strep). 100 ng/well of each plasmid was transfected. 24 hours after transfection, the complete DMEM media was removed, and 100 μl of Opti-MEM reduced serum was added. The assay used the Nano-Glo Live Cell Assay System (#N2011, Promega). The Nano-Glo Live cell reagent is reconstituted by combining 1 volume of Live cell substrate with 19 volumes of LCS dilution buffer. 25 μl of Live cell reagent is added to each plate and gently mixed by hand, followed by readings on the Spark (Tecan) microplate reader.

### CRISPR/Cas9 gene editing

HiBiT tag was edited into U2OS cells by electroporating the RNP complex using the Neon Transfection System (Invitrogen). The RNP complex consists of Alt-R^TM^ S.p. Cas9 V3 enzyme (IDT), crRNA (IDT) targeting PVT1 Ex9 or Ex2 and tracrRNA (IDT). To form gRNAs designed crRNAs, tracrRNA is complexed by combining 100 mM of each in a tube and heating to 95°C for 5 minutes. To prepare the RNP complex, 36 pmol of each gRNA, and 60 pmol of Cas9 enzyme are added to a tube with 1X PBS to a total volume of 3ml and incubated at room temperature for 10 minutes. 100,000 U2OS cells are suspended in 6 ml of R buffer is added to the test tube containing 3 ml of the RNP complex and 100 pmol of HDR template. R buffer is added to bring the volume to 15 ml. The transfection mix is pipetted into 10 ml Neon Tips using the Neon Pipette carefully, avoiding bubbles. Transfection was performed according to the manufacturer’s specifications for U2OS cells. After electroporation, cells were transferred into 24-well plates containing warm media. Negative control crRNA (IDT) was used as control. After 72 hours cells were enriched and tested for positive cells using the Nano-Glo HiBiT lytic detection assay (Promega). Positive wells were further enriched to obtain single-cell clones. The Nano-Glo HiBiT lytic detection assay identified successful clones and the in-frame knock-in was confirmed through Sanger sequencing.

### Nano-Glo HiBiT Lytic Detection Assay

To prepare Nano-Glo Lytic Reagent, 1:100 LgBiT protein and 1:50 Nano-Glo HiBiT Lytic Substrate were added to room temperature Nano-Glo HiBiT Lytic Buffer. After mixing the reagents, equal volumes of the Nano-Glo HiBiT Lytic reagent as the media were added to HiBiT edited cells in a 96-well plate. The plate was placed on an orbital shaker for 10 minutes and then incubated at room temperature for equilibration after the mixing. Luminescence was recorded on a Tecan microplate reader. Unedited cells were used as a negative control.

### Immunofluorescence Imaging

To study the expression and distribution of HNB in cancer cells. We imaged HNB-HiBiT edited cells using Anti-HiBiT antibody and Anti-mouse Alexa-Fluor 488 conjugate antibody to visualize HNB in HNB-HiBiT U2OS cells. Cells were fixed with 4% PFA for 15 mins, and permeabilized with 0.2% Triton X-100 (1X PBS, 0.2% Triton X-100). They were then washed 3 times with 1X PBS and incubated with 3% BSA in 1X PBS at room temperature for 60 minutes. Anti-HiBiT antibody (#N7200, Promega) is added at 1:250 dilution and incubated overnight at 4°C on an orbital shaker. After 16 hours of incubation the primary anti-body is removed, and the cells are washed 3 times with 0.2% Triton X-100 (1X PBS, 0.2% Triton X-100). 1:1000 dilution Anti-Mouse Alexa-Fluor488 Conjugate (#4408, Cell Signaling Technologies) was added to the cells and incubated for 60 minutes at room temperature on an orbital shaker. The antibody is washed 3 times with 1X PBS and mounted on a glass slide using ProLong Gold antifade reagent with Dapi (#P36931, Invitrogen). Images were taken on the Nikon Super Resolution Microscope and analyzed on Nikon Elements.

### Quantification and Statistical Analysis

Statistical analyses for figures were performed using GraphPad Prism software (San Diego, CA). Data are presented as the mean ± standard deviation unless otherwise specified. For quantitative PCR experiments, Gaussian distribution was assumed, and a student t-test (2-tailed unpaired) was used to determine statistical significance. Significant differences between groups were determined using a student t-test (2-tailed unpaired). All experiments were performed at least 3 independent times unless otherwise noted. The significance level for statistical testing was set at P < 0.05. No statistical method was used to predetermine any other sample sizes.

### Bioinformatic analysis (Gene Set Enrichment Analysis)

RNAseq processing: Raw reads were mapped to the hg19 human reference genome using the STAR aligner (v.2.5.0a) 66. PCR-duplicated reads and low mapping quality reads (MQ score below 20) were removed using Picard tools and SAMtools, respectively. Aligned reads were counted using htseq-count (v.0.6.1p1) 69 and normalized according to size factors method from DESeq2 70. We used fold change ≥2 and FDR <0.05 as the determination of differentially expressed genes (DEG). Statistical analysis and plots were performed inside R environment version 3.1.0. upload files)

### Identification of Honeybadger and Firefox peptides using PepQuery2

The Honeybadger and Firefox peptides were identified using PepQuery2 which matches the peptides to publicly accessible mass spectra data from various cancer cohorts and healthy tissues. To identify the peptide the Honeybadger sequence was searched against GTEx_32_Tissues, CPTAC-CCLE, CPTAC-PDA and CPTAC-GBM datasets. The parameters set were, novel peptide/protein, the target event was set to protein sequence, gencode_v38_human was used as a reference database and the scoring was done using Hyperscore. A total of 37 PSMs were identified with confidence from 1 healthy tissue and 4 cancer datasets. The representative mass spectra of the Honeybadger peptide TQLGAVK identifies it in a GTEx_32_tissue sample.

The Firefox protein sequence was searched against Deep_32_healthy_tissue and CCLE database using the same parameters described above. A total of 2 PSMs were identified with confidence from the healthy tissue and cancer database. The representative mass spectra of the Firefox peptide MHVPSGAQLGLRPDLLAR identifies it in a Deep_32_healthy_tissue sample

### Computational analysis of KRAS-HNB interaction

The AlphaFold Server (https://alphafoldserver.com/) was used to assess the interaction between KRAS and HNB in a 1:1 ratio with randomized seeding. The results were visualized using UCSF ChimeraX ^64^

### Whole Genome Sequencing

Genomic DNA was sheared on a Covaris S2 (Covaris Inc.) and libraries were made using the NEBNext Ultra II DNA Library Prep Kit for Illumina (NEB, Inc.). Indexed libraries were pooled, and paired end sequenced (2x75bp) on an Illumina NextSeq 500 sequencer. Read data was processed in BaseSpace (basespace.illumina.com). Reads were aligned to Homo sapiens genome (hg19) using BWA aligner version 0.7.13 (https://github.com/lh3/bwa) with default settings.

### Data Availability

The data that support the findings of this study are available from the corresponding author upon request and will be fully access before final publication.

### Inclusion & Ethics Statement

#### Human Participants

All studies involving human tumor samples were conducted in compliance with international guidelines and were approved by the Institutional Review Board of the Sick Kids Hospital, Toronto, Canada.

#### Animal Studies

All animal procedures were conducted in accordance with the guidelines established by the Institutional Animal Care and Use Committee of Sanford Burnham Prebys Medical Discovery Institute. Efforts were made to minimize animal suffering and to reduce the number of animals used as per the guidelines of AAALAC International and a Multiple Project Assurance A3053-1 is on file in the OLAW, DHHS.

#### Inclusion and Diversity

We have actively ensured that the study’s design, conduct, and reporting were free from bias, regardless of gender, age, ethnicity, socio-economic status, or disability of the participants. We believe in and adhere to principles of equity and inclusivity in research.

## Acknowledgements

We thank the members of the Wechsler-Reya and Taylor laboratories for helpful discussions, the animal core facility staff for help with mouse colony. We thank Drs Vineet Bafna and Howard Chang for their insight and suggestions. We also thank Dr. Paul Mischel for help and suggetions with the dual FISH assay. A.B. was supported by NIH/NCI (1R01CA200643-01A1), Department of Defense/PCRP (PC160884), American Cancer Society Research Scholar Grant (125627-RSG-14-074-01-TBG), Tobacco-Related Disease Research Program (T33IR6727) and Curebound (CURE22).

## Author Contributions

A.T and A.B. conceived the project. A.T, K.T, U.P, A.V., J.T.L, S.W, M.B.M., A.S., K.V., J.F, B.H., A.S., L.D, performed experiments. O.S., K.B., Q.T., T.N., T.M., L.H, B.J., O.C., J.L. analyzed data. B.K, S.M.D., Y.M., L.S., L.C., V.B., P.S.M., H.S., M.I.H., A.J.D., R.W.R., M.D.T. and A.B. guided data analysis and provided feedback on experimental design. A.T, R.W.R. and A.B. wrote the manuscript with input from all authors.

## Materials & Correspondence

Correspondence and requests for materials should be addressed to Anindya Bagchi

**Primer sequences.** All primers used for library amplification or RT-qPCR are listed.

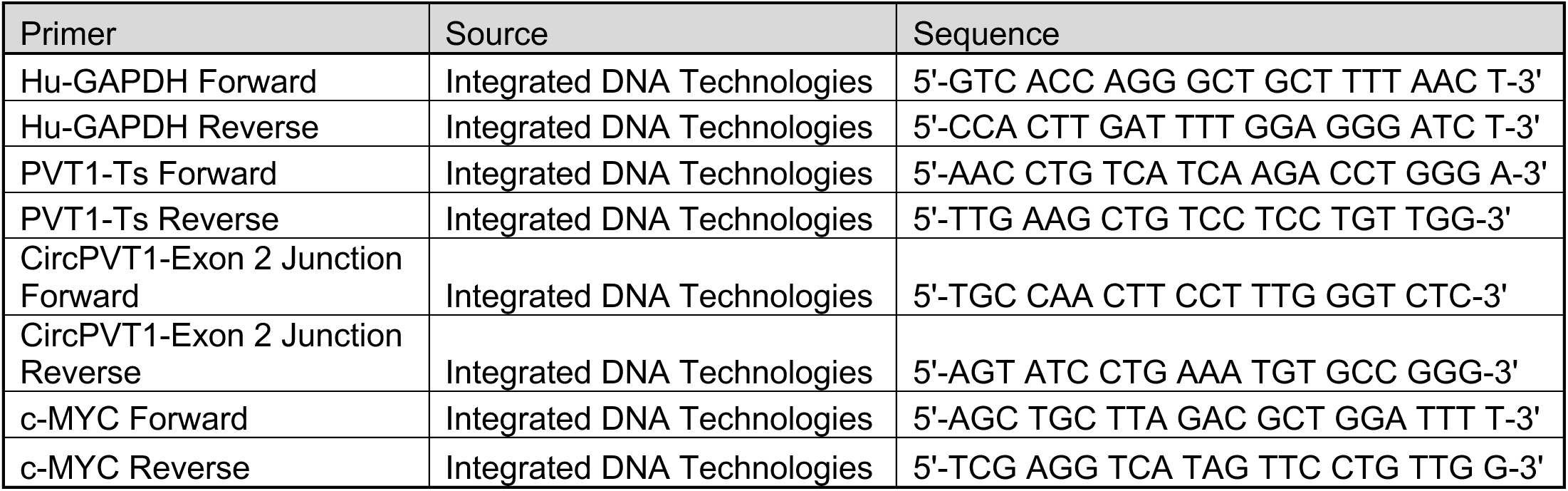

**si-RNA/sh-RNA sequences.**

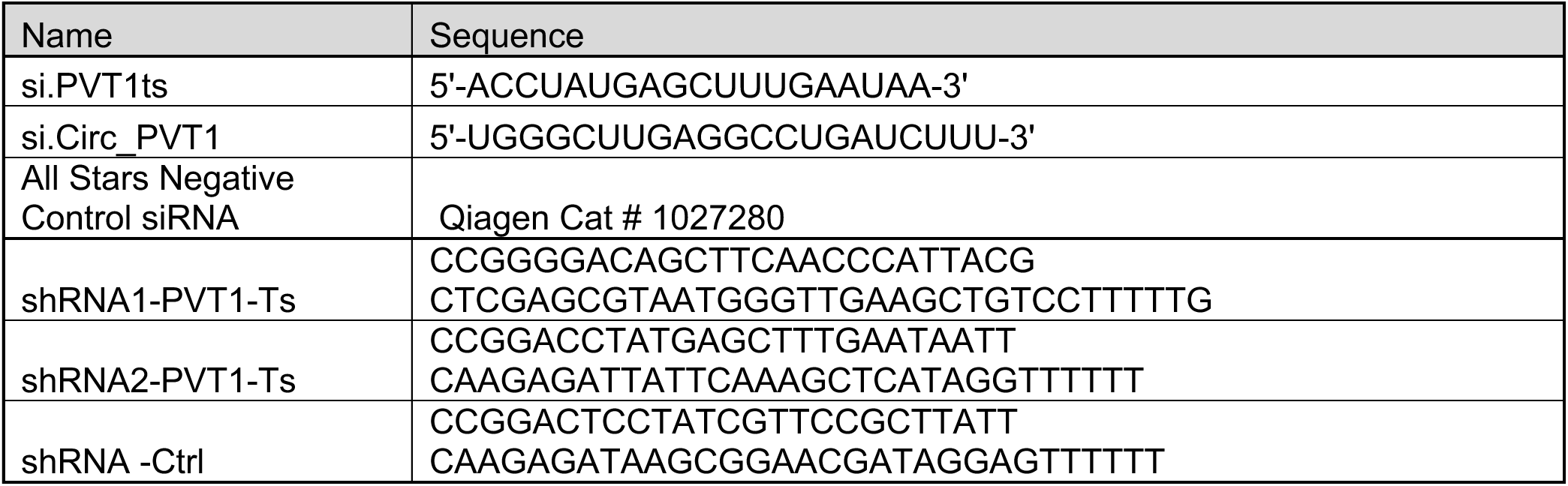

---

**Supplementary Figure 1:**
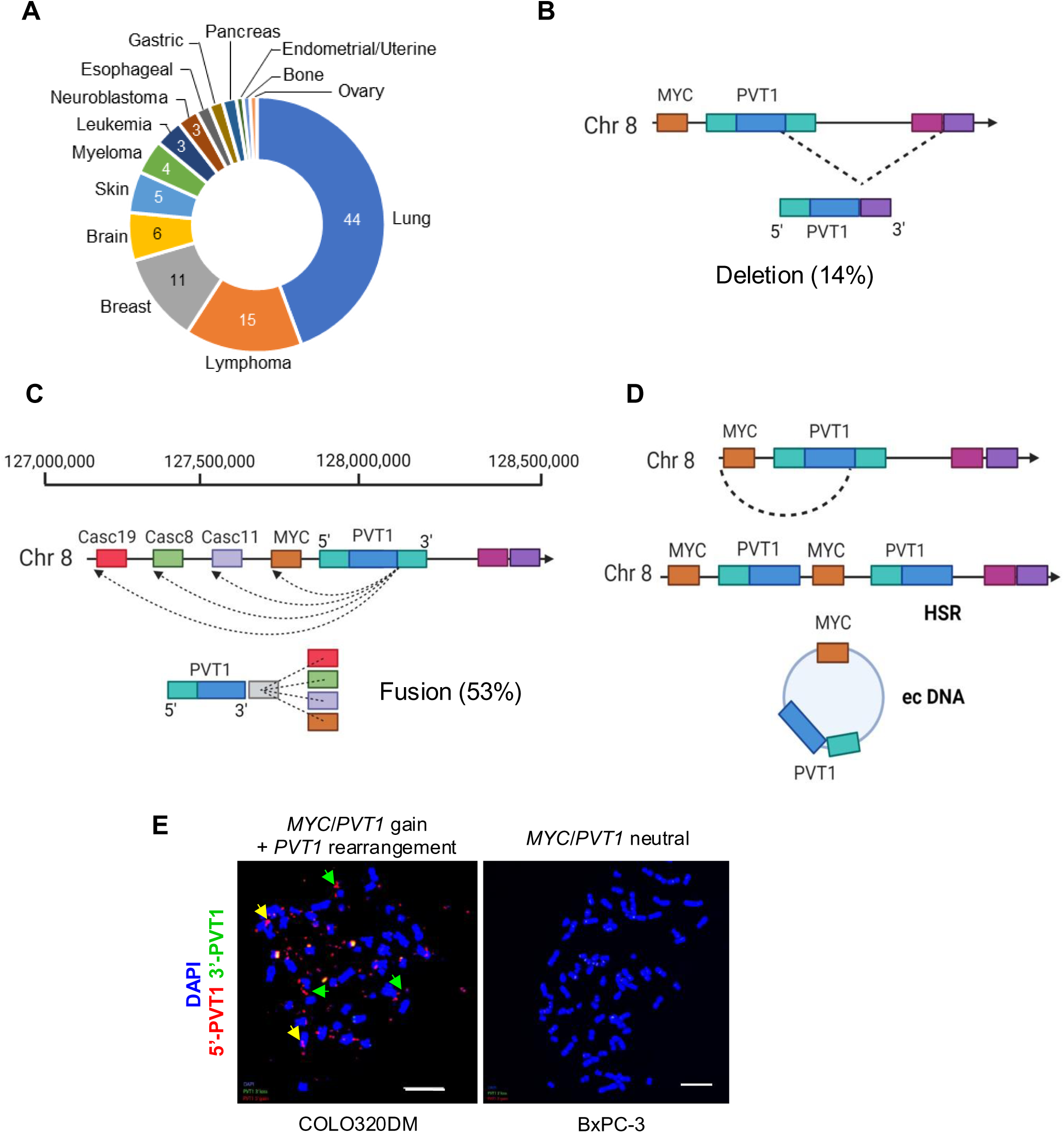
PVT1 fusions in cancers (A) Donut plot showing the distribution of PVT1 fusion-positive cancers in the CCLE dataset. The slice for each cancer type is proportional to the number of cell lines for that cancer type. A total of 115 cell lines were positive for PVT1 fusion (n=115). (B) 14% of intrachromosomal PVT1 fusions harbor deletion of the PVT1 gene. (C, D) 53% of intrachromosomal rearrangements fuse 5’-PVT1 to a gene partner on 8q24 located upstream of PVT1, such as CASC11, CASC8, or MYC, suggesting that such fusions may generate extrachromosomal DNA (ecDNA) and/or double minute chromosomes (HSRs). (E) Representative dual probe (5’-PVT1/3’PVT1) FISH images of COLO320DM (MYC/PVT1 gain + PVT1 rearrangement) and BxPC-3 (MYC/PVT1 neutral) cell lines. COLO320DM showed 5’ enrichment of PVT1 on ecDNA and HSR, as shown by the green and yellow arrows, respectively. BxPC-3 cells were used as control. Scale bars, 10 µm.

**Supplementary Figure 2:**
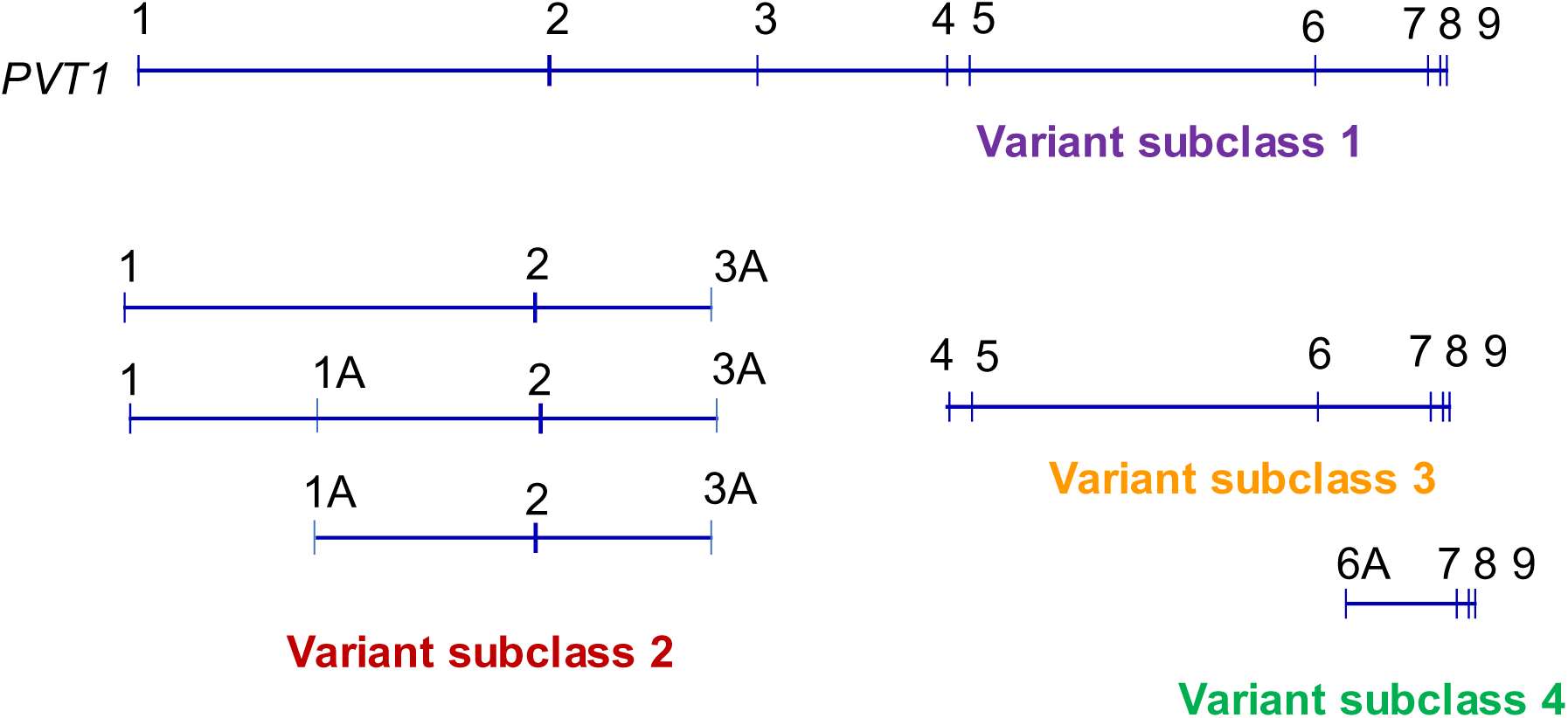
Broad subcategorization of 183 PVT1 transcript variants (ENSG00000249859) The primary PVT1 transcript contains nine exons (1-9). 183 PVT1 transcript variants were identified. These variants were broadly categorized into four subclasses: Variant subclass 1 (18 variants): Contains full-length PVT1. Variant subclass 2 (67 variants): Contains exons 1, 2, and 3A, and similar iterations spanning 5’-PVT1. Variant subclass 3 (86 variants): Contains exons 4A-9 (and their different sub-iterations). Variant subclass 4 (22 variants): Contains exons 6A-9. (and their different sub-iterations). The distribution of variants across subclasses is shown in Supplementary Table 1.

**Supplementary Figure 3:**
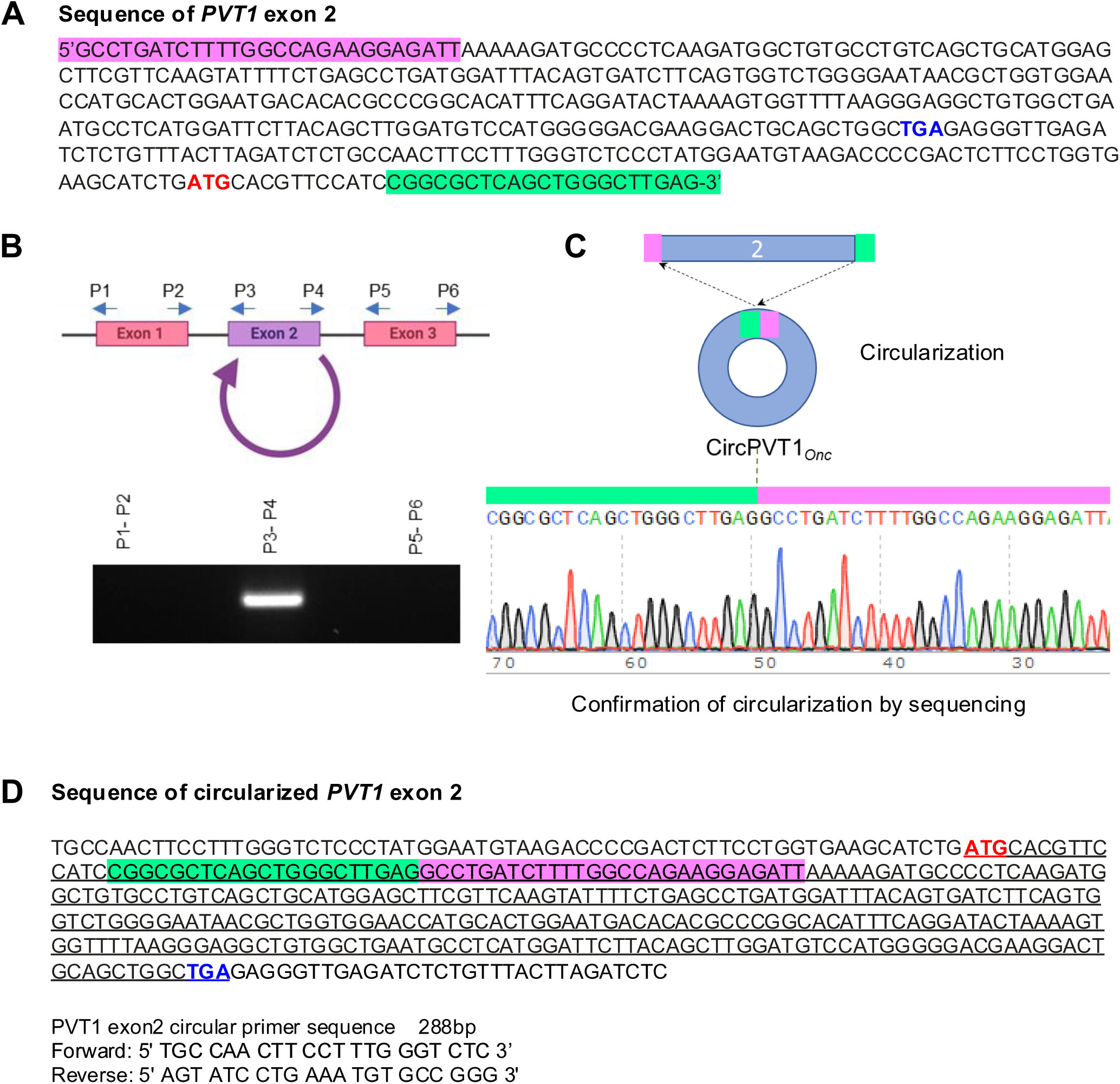
PVT1 exon 2 forms Circular RNA (A) Nucleotide sequence of PVT1 exon 2. The 5’ and 3’ ends of exon 2 are highlighted by pink and green color, respectively. The translation start and stop codons are highlighted. (B) Schematic representation of divergent primer pairs (P1/P2, P3/P4, and P5/P6) for respective exons to identify circularization. Agarose gel electrophoresis of amplified product from cDNA (prepared by random hexamer oligonucleotides) using the described primer pairs. (C) Schematics of exon 2 circularization and confirmation of junction sequence (joining of 5’/3’ ends) by sequencing of amplified product (P3/P4 primer pair). (D) Nucleotide sequence of circularized exon 2 and resulting new ORF is underlined with start and stop codon highlighted in red and blue color, respectively.

**Supplementary Figure 4:**
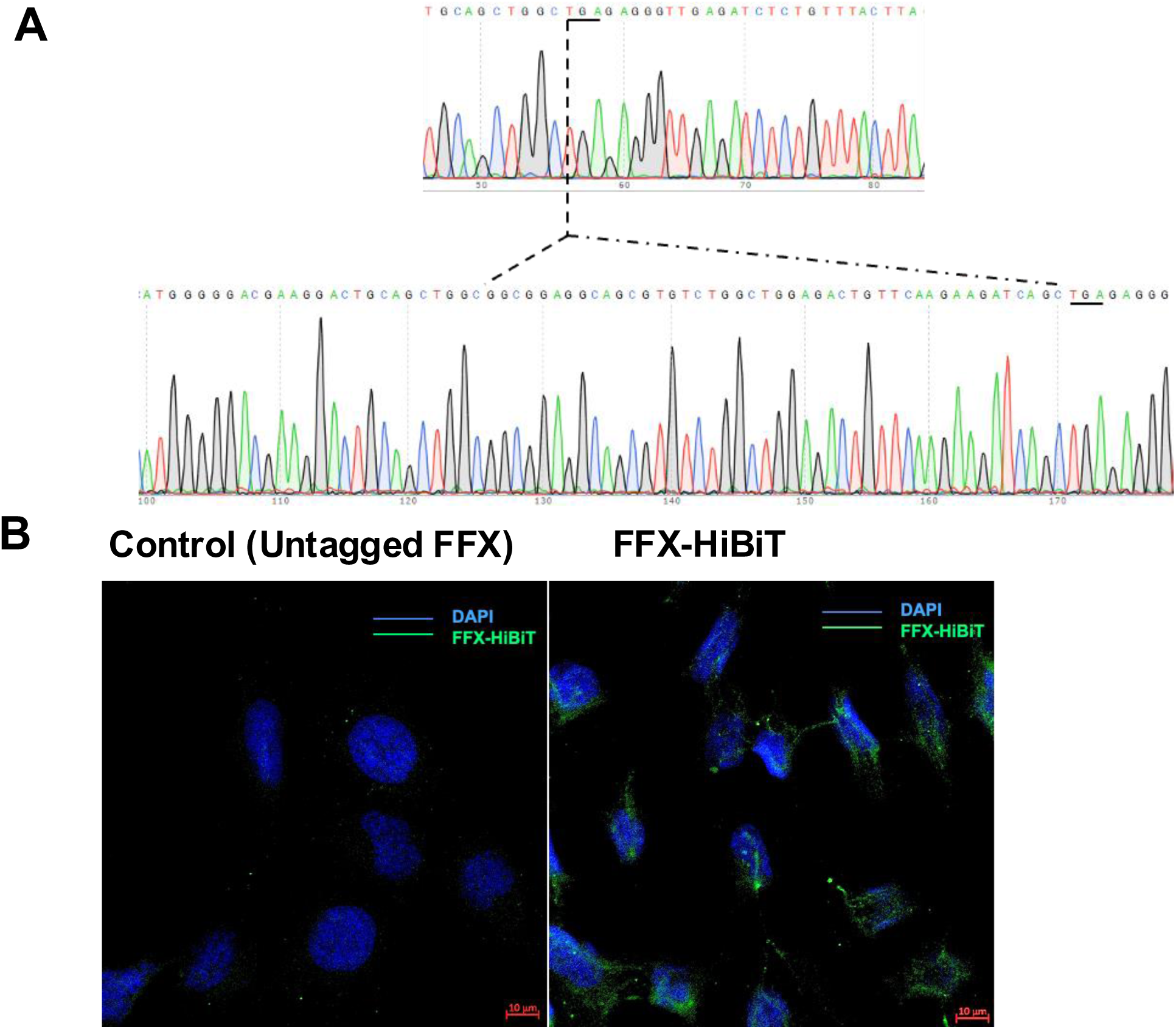
Endogenous tagging of FFX with HiBiT. (A) Sequence chromatograms of HiBiT sequence fused to endogenous FFX (immediately upstream of native stop codon to generate C-terminal protein fusion) using CRISPR/Cas9 knock-in. The nucleotide sequences of the HiBiT tag are also shown. (B) Immunofluorescent detection of HiBiT-tagged FFX in CRISPR-edited cell clones using the anti-HiBiT monoclonal antibody (green) and DAPI (blue). Scale bars, 10 µm.

**Supplementary Figure 5:**
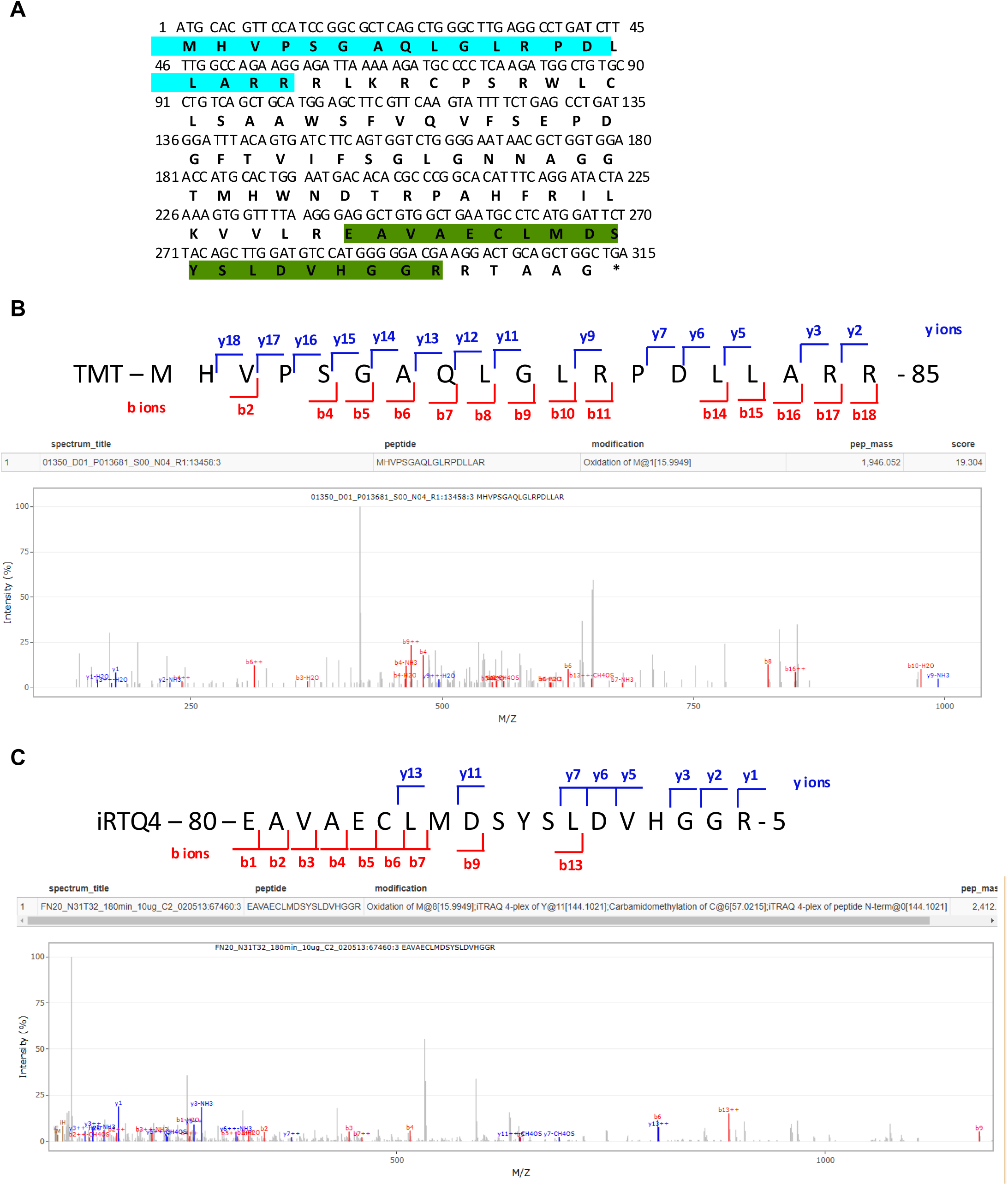
Identification of FFX peptide sequence in human proteomic databases. **(A)** FFX sequence with the highlighted regions corresponding to the peptides identified in the human proteomics database. **(B, C)** Representative Mass spectra image of FFX peptides MHVPSGAQLGRPDLLAR and EAVAECLMDSYSLDVHGGR identified from the human proteomics database. The y-axis represents relative ion abundance, while the x-axis represents the m/z ratio.

**Supplementary Figure 6:**
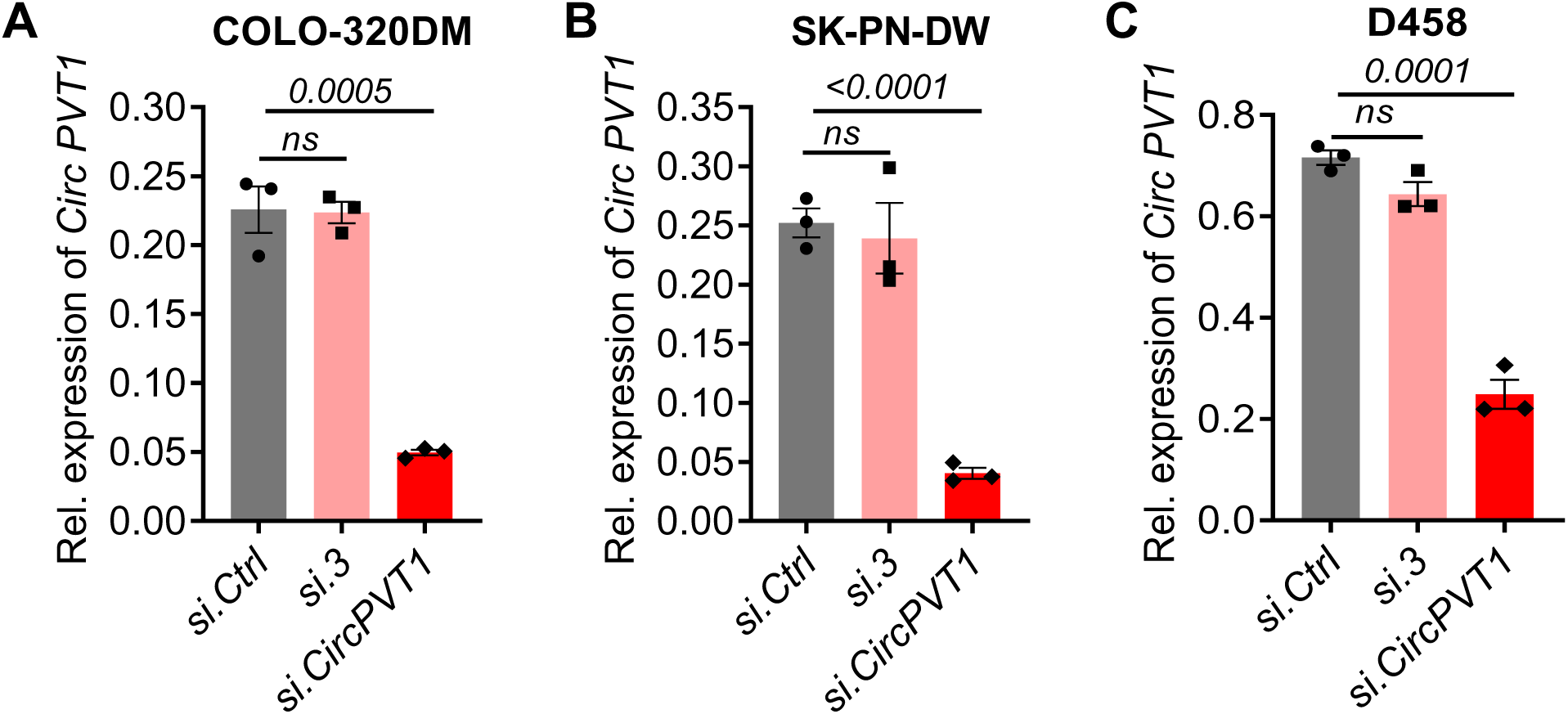
siRNA mediated knockdown of CircPVT1. qRT-PCR of CircPVT1 transcript from (A) COLO-320DM, (B) SK-PN-DW and (C) D458 cells after transfection with si.Ctrl (control siRNA), si.3 (siRNA targeting linear PVT1), and si.CircPVT1 (siRNA targeting CircPVT1). (n=3, P values were determined by unpaired t test.)

**Supplementary Figure 7:**
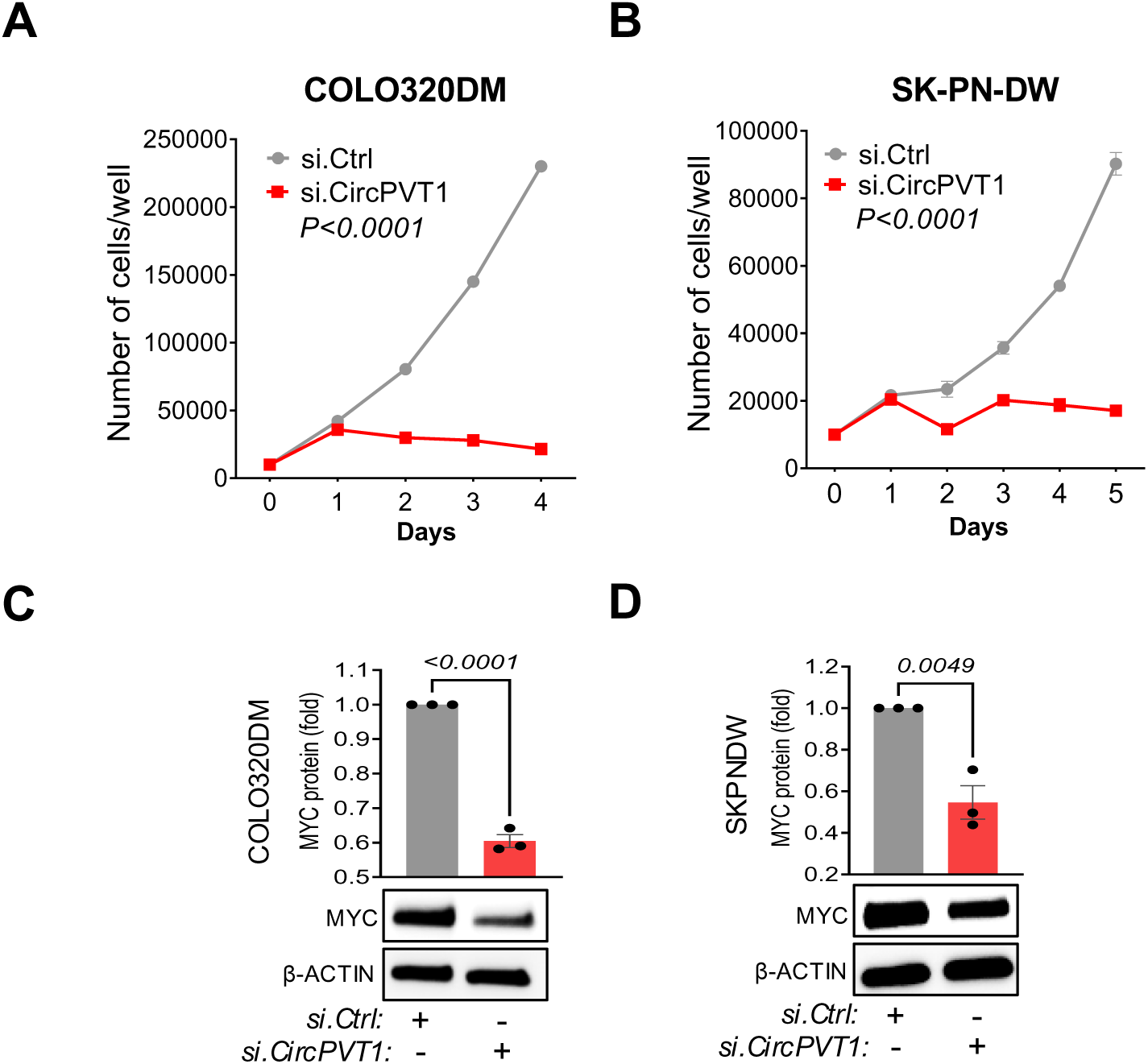
Cell proliferation assay of COLO320DM (A) and SK-PN-DW(B) after transfection with si.CircPVT1 or si.Ctrl (n=3, P value by two-way ANOVA). (C-D) Western blot analysis of MYC expression in COLO320DM and SK-PN-DW transfected with si.Ctrl or si.CircPVT1. β ACTIN was used as loading control (n=3, P value by unpaired t test).

**Supplementary Figure 8:**
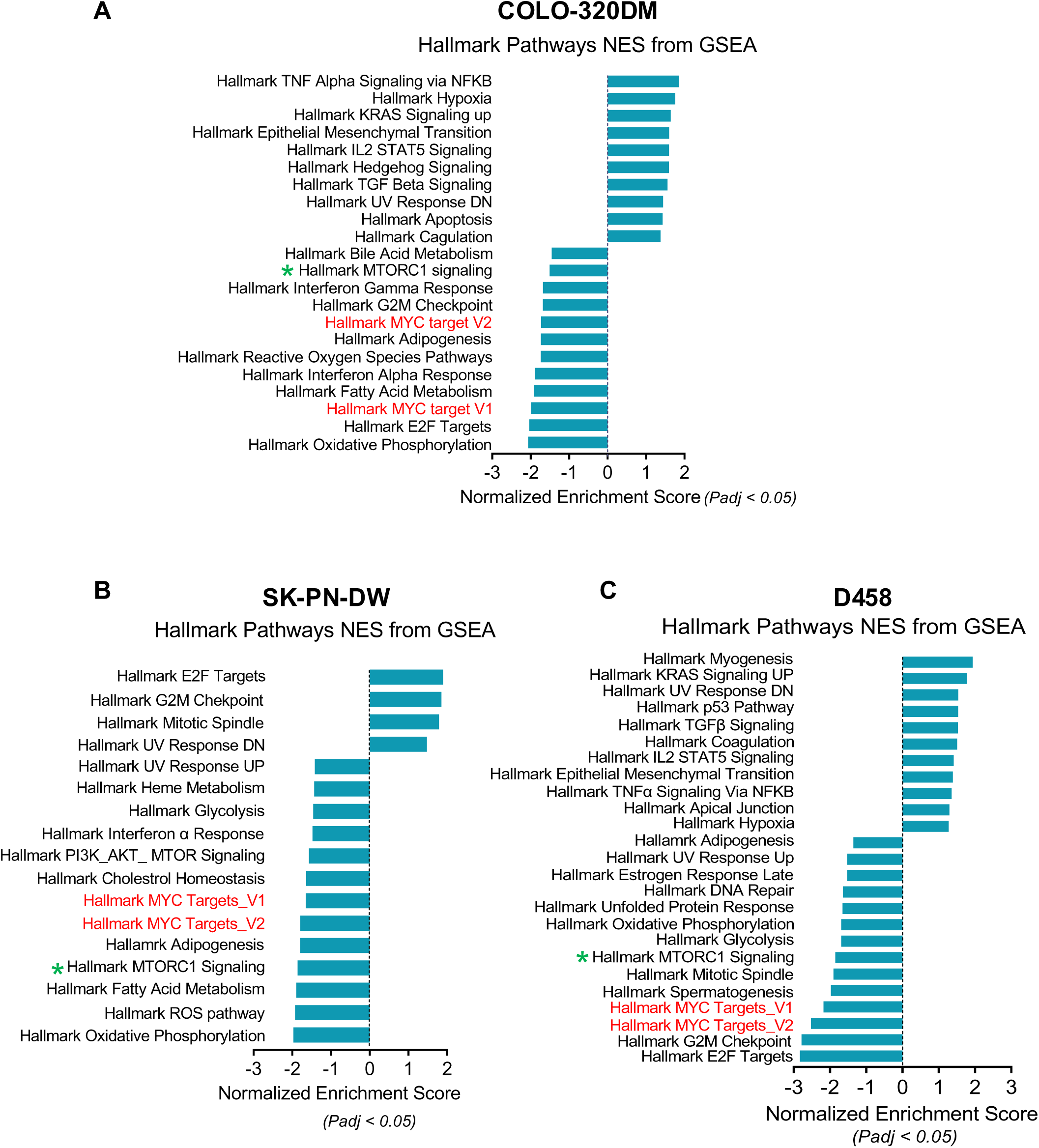
Global hallmark pathway analysis of RNA Seq data from si.CircPVT1 vs si.Ctrl treated cells. Gene set enrichment analysis (GSEA) of differentially expressed protein pathways in (A) COLO-320DM, (B) SK-PN-DW, and (C) D458 cells after knocking down CircPVT1 (n=3). The X and Y axes represent normalized enrichment scores (NES) and the total list of GSEA hallmark upregulated and downregulated categories, with significantly enriched terms (p-adjust < 0.05). The hallmark MYC targets V1 and V2 (in red) and MTORC1 (in green asterisk) are highlighted

**Supplementary Figure 9:**
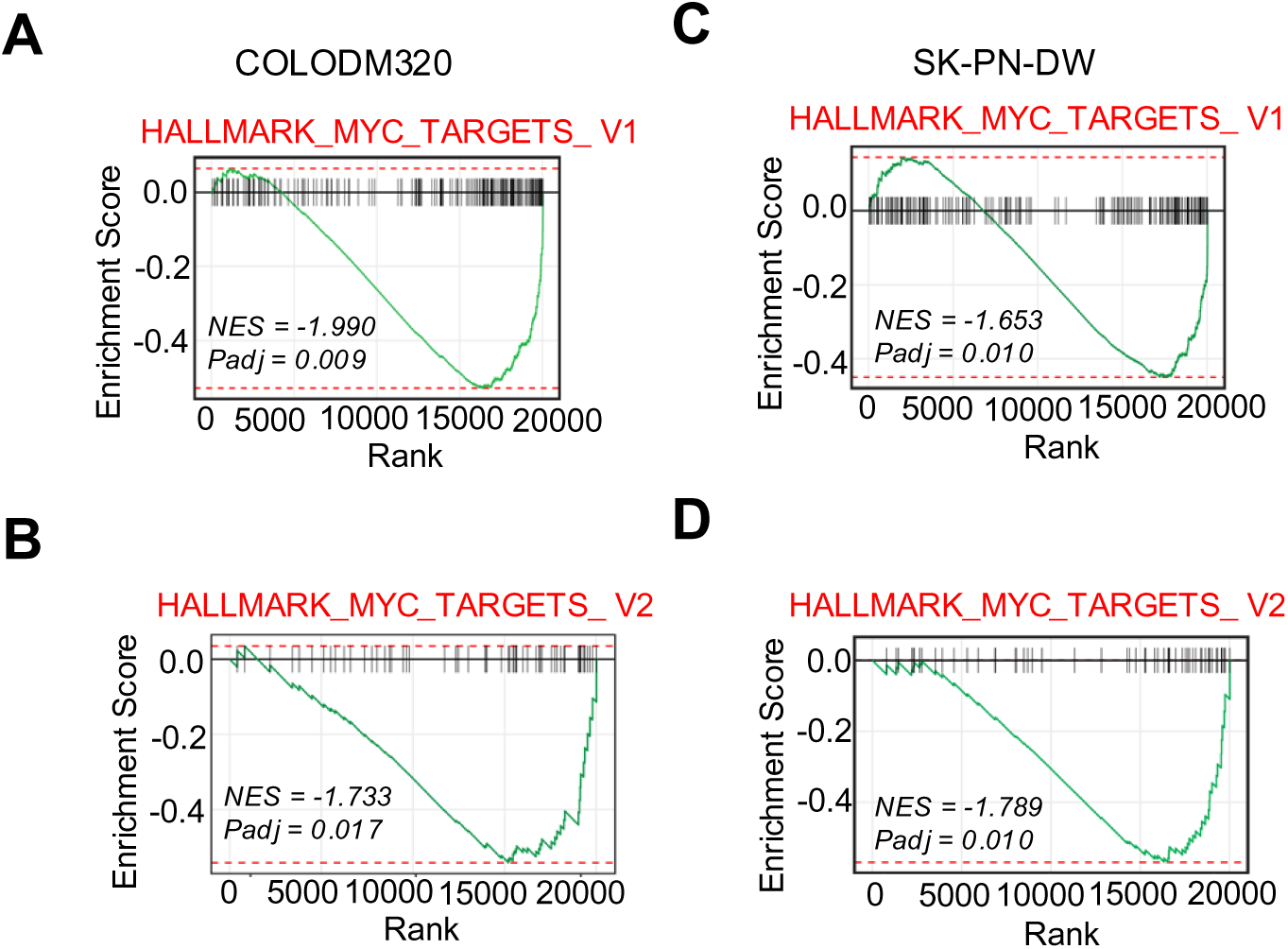
Gene set enrichment analysis scores (GSEA) analysis for MYC targets genes in COLODM320 (A,B) and SK-PN-DW (C,D) treated with si.CircPVT1. si.Ctrl was used as control. n=3, for each treatment condition.

**Supplementary Figure 10:**
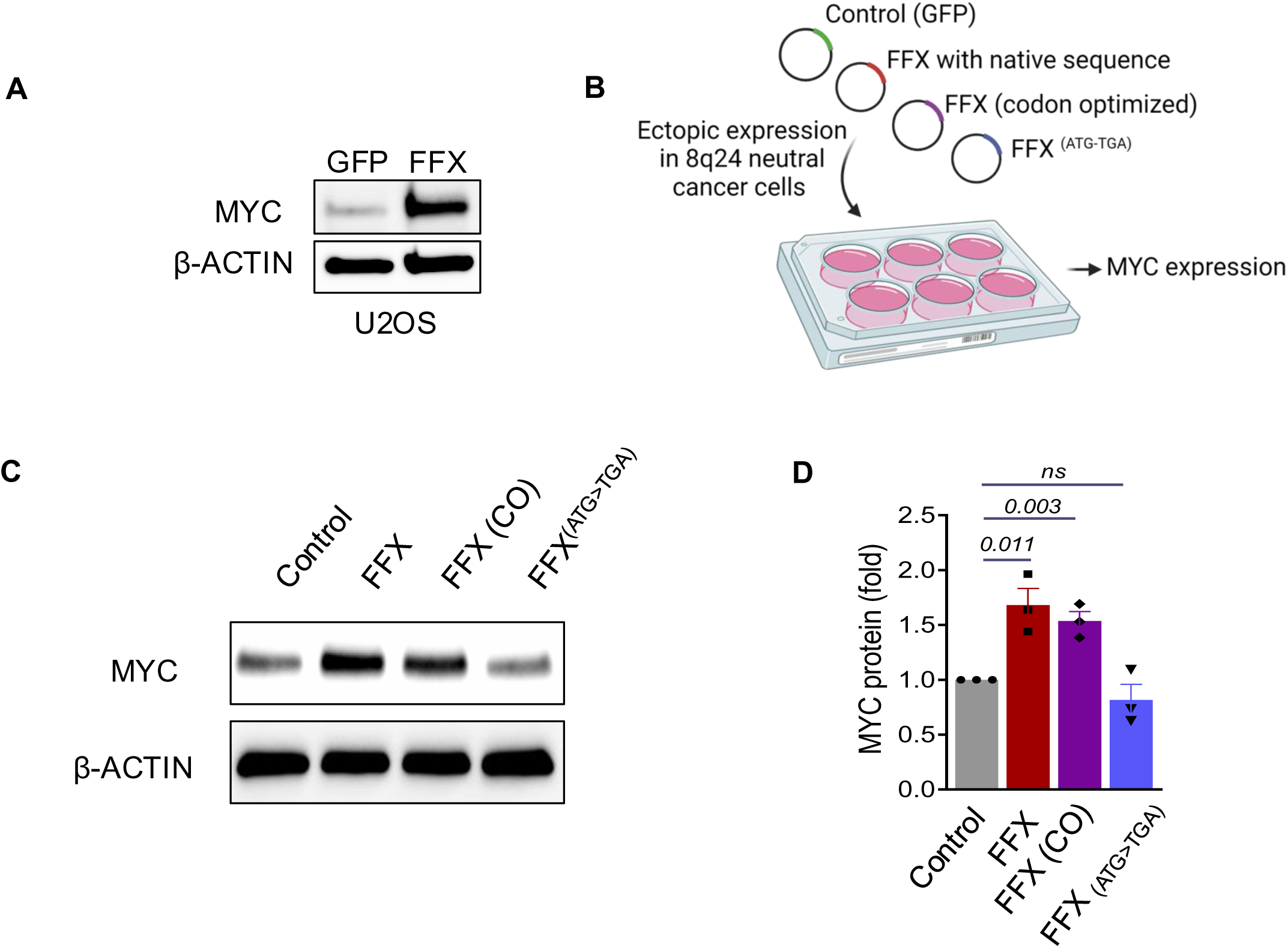
FFX protein augments MYC expression **(A)** Western blot analysis of MYC expression in U2OS cells ectopically transduced with constructs expressing GFP (control) or FFX. **(B)** Schematic of assay for MYC expression in U2OS cells ectopically transduced with constructs expressing GFP (control), FFX (with native sequence), codon optimized FFX (FFX(CO)), or mutant FFX in which the start codon ATG is mutated to TGA (FFX^ATG>TGA^). **(C&D)** Western blot analysis and quantification of MYC expression in U2OS cells transduced with GFP (control), FFX, FFX(CO), and FFX ^ATG>TGA^. (n=3, p value by unpaired t test).

**Supplementary Figure 11:**
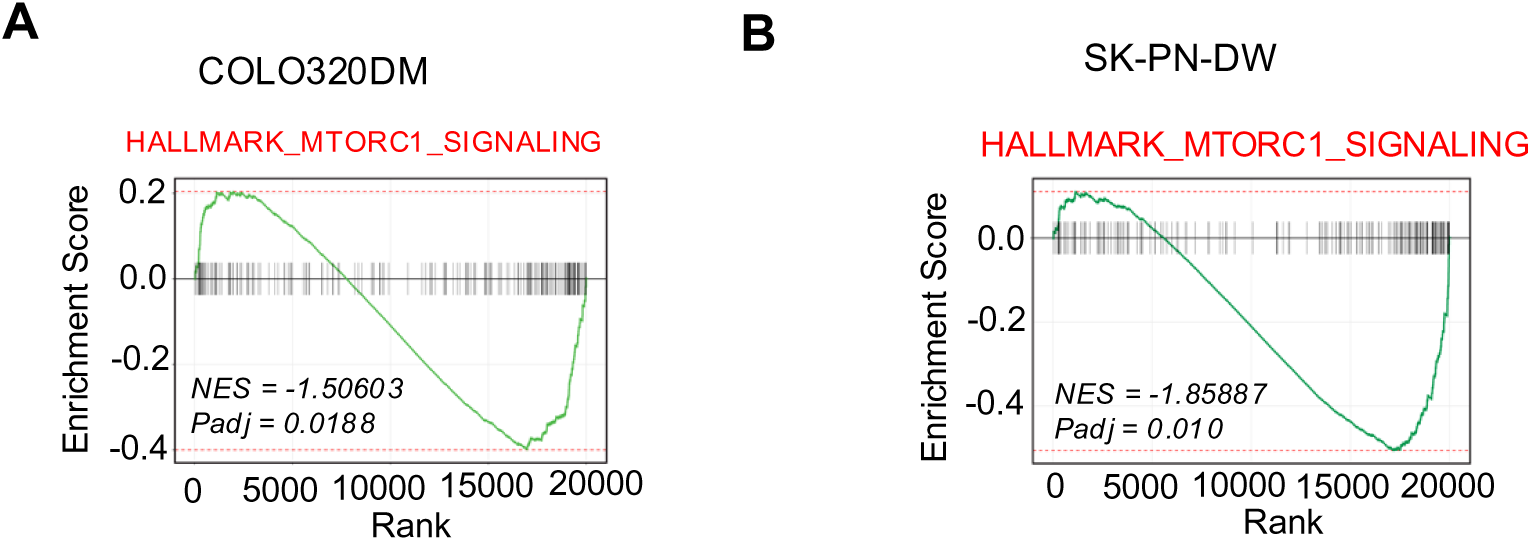
Gene set enrichment analysis scores (GSEA) analysis of genes involved in MTORC1 signaling in COLO320DM (A) and SK-PN-DW (B) treated with si.CircPVT1. si.Ctrl was used as control. n=3, for each treatment condition.

**Supplementary Figure 12:**
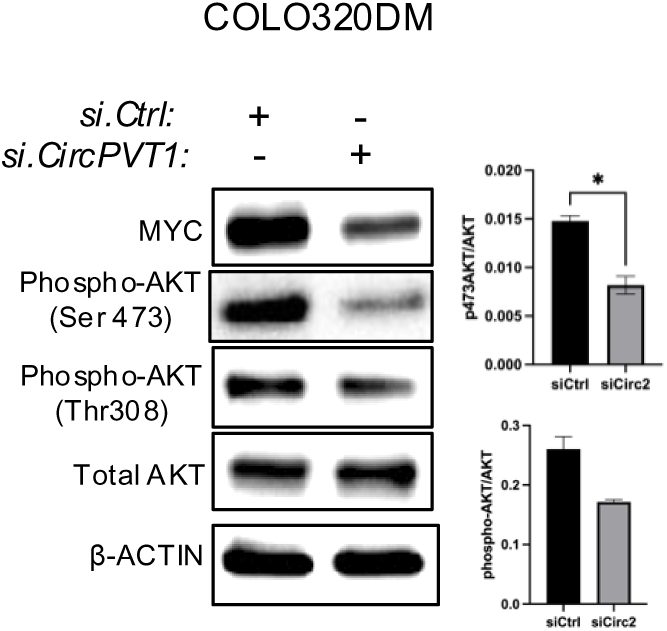
Western blot and quantitative analysis of MYC, phospho-AKT (Ser 473 and Thr308), total AKT expression in D458 cells transduced with si.CircPVT1 and si.Ctrl. β ACTIN was used as a control. (p-values obtained by unpaired t-test, n=3, for each treatment condition)

**Supplementary Figure 13:**
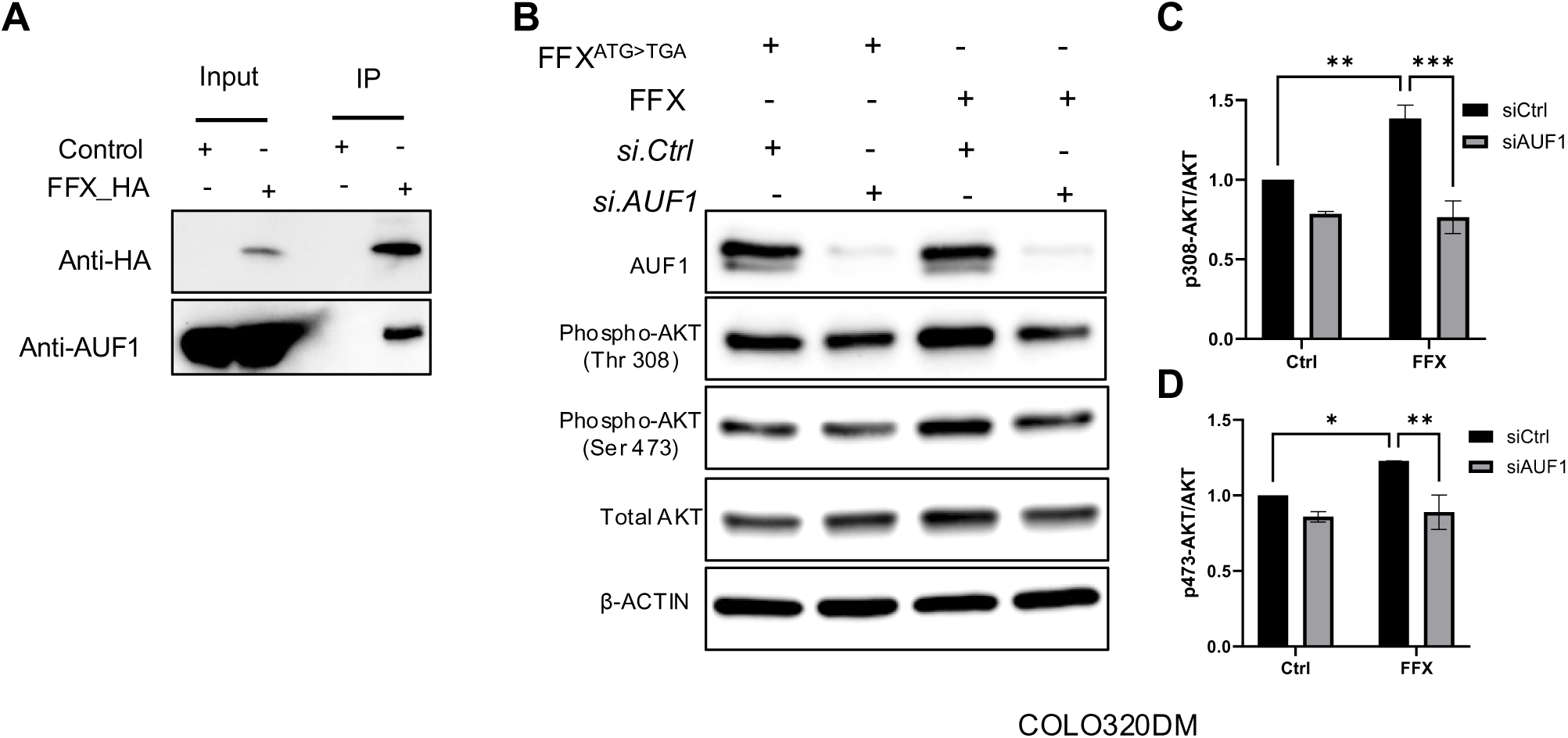
**A.** Western blot analysis of Co-IP experiments from U2OS transfected with FFX (control) or HA tagged FFX (FFX_HA) with indicated antibodies**. B.** Western blot analysis of AUF1, phospho-AKT (Thr308 and Ser 473), total AKT expression in COLO320DM cells transduced with FFX^ATG>TGA^ (Control) and FFX and treated with si.AUF1 and si.Ctrl. β ACTIN was used as a control. (**C,D**) Quantitative analysis of phospho-AKT (Thr308 and Ser 473) expression in COLO320DM cells transduced with FFX^ATG>TGA^ (Control) and FFX and treated with si.AUF1 and si.Ctrl (p-values obtained by unpaired t-test, n=3, for each treatment condition).

**Supplementary Figure 14:**
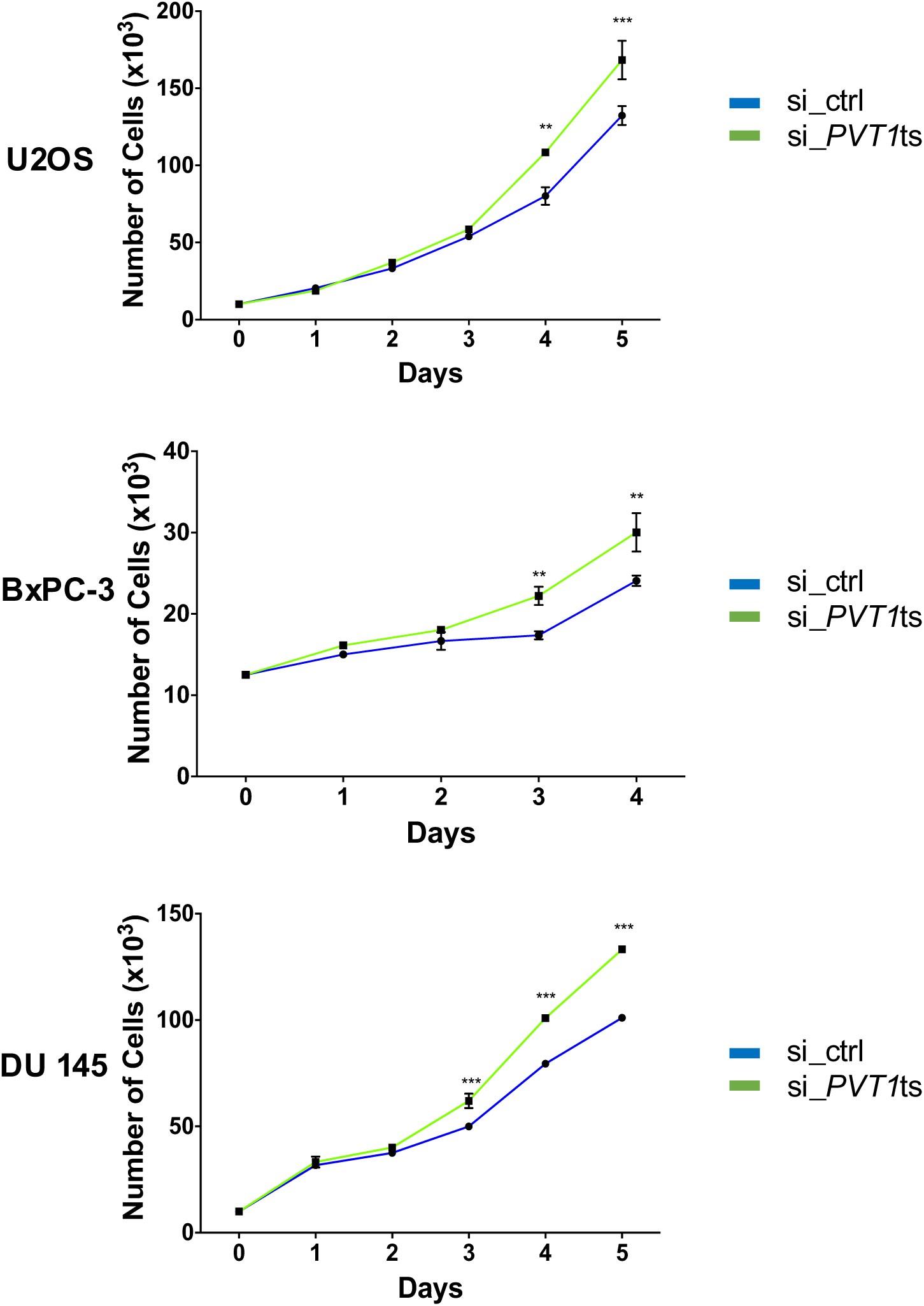
Analysis of Cell Proliferation in MYC/PVT1 Neutral Cell Lines (U2OS, BxPC-3, DU-145) following Transfection with si_PVT1ts or si_ctrl: Each experiment was performed in triplicate (n=3), and statistical significance was determined using two-way ANOVA (P values indicated)

**Supplementary Figure 15:**
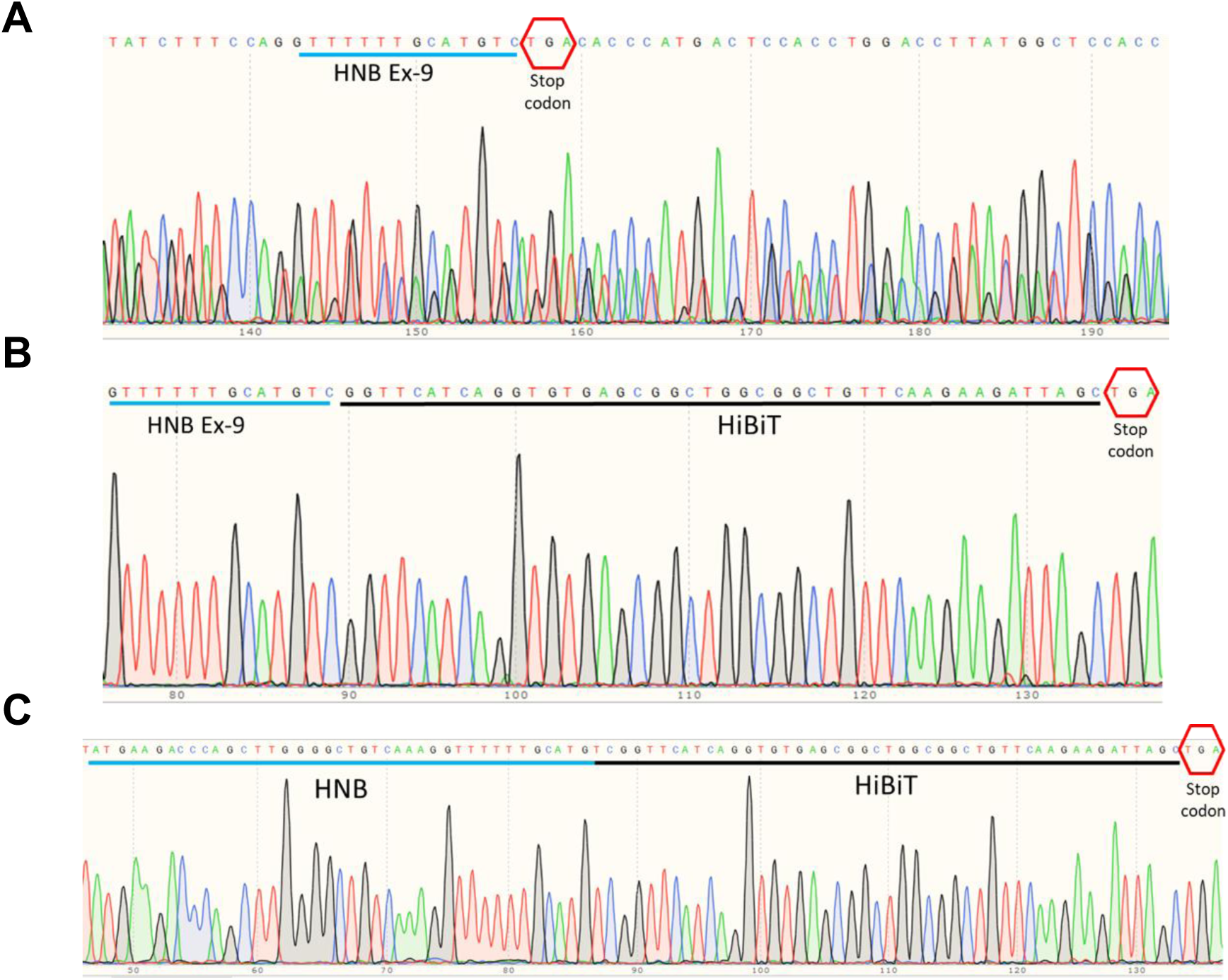
CRISPR/Cas9 mediated integration of HiBiT tag at HNB’s C-terminus in U2OS cells: Genomic sequence chromatograms comparing native HNB (untagged, **A**) and HiBiT-tagged HNB (**B**). The chromatograms display bioluminescent-negative control cells (A) and bioluminescent-positive HiBiT-tagged U2OS cells (B), confirming the precise knock-in of the HiBiT tag before the stop codon in PVT1 Exon9 of HNB. (C) Sequence chromatograms of HNB-HiBiT cDNA from the HiBiT-positive U2OS cell line, verifying the expression of HNB-HiBiT mRNA

**Supplementary Figure 16:**
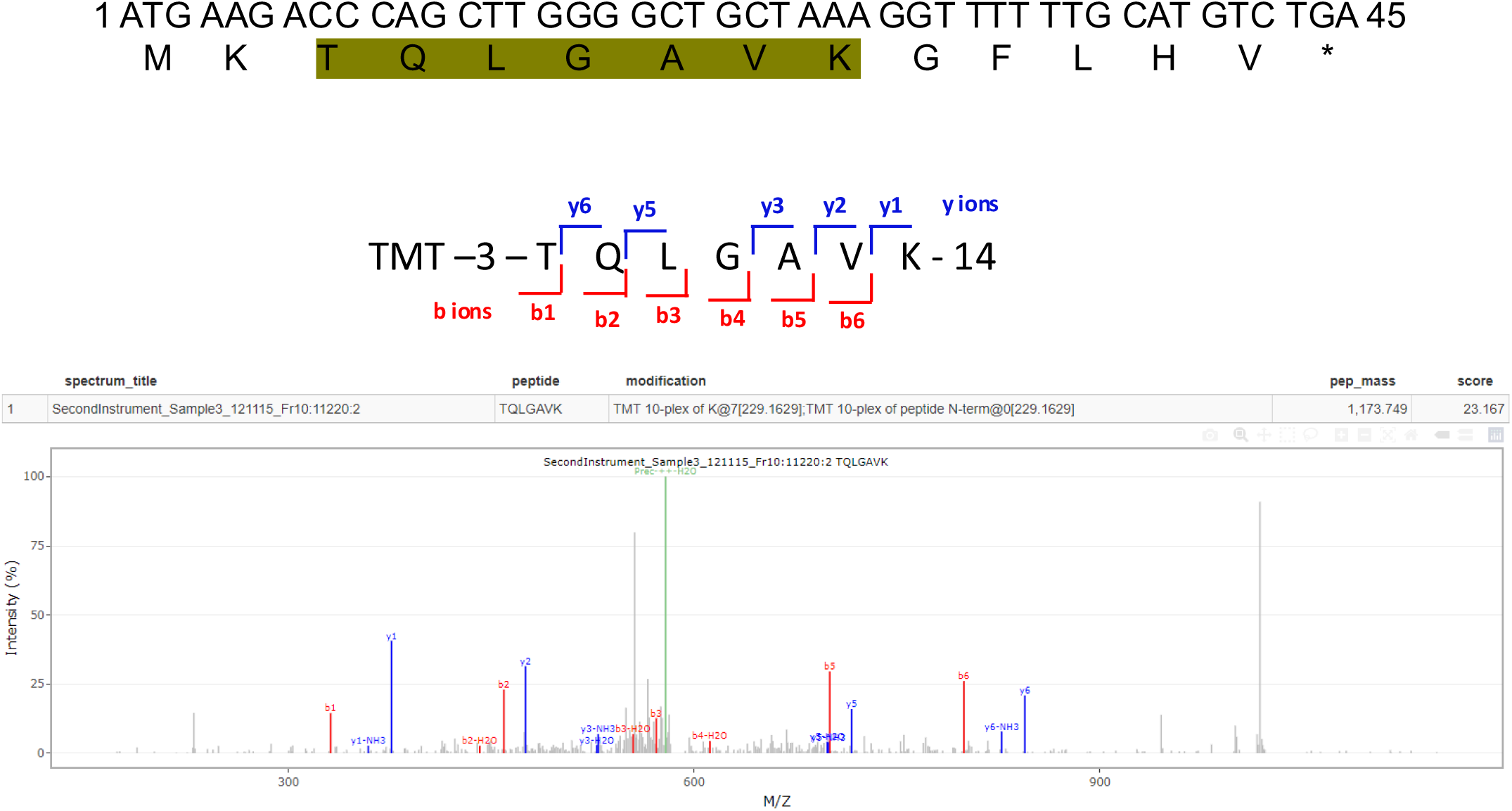
Identification of HNB peptide sequence in human proteomic databases. **(A)** HNB peptide sequence with the highlighted region corresponding to the peptide identified in the human proteomic database. **(B)** Representative MS/MS spectra image of HNB peptide TQLGAVK identified from GTEx_32_Tissues_Proteome_PXD016999. The y-axis represents relative ion abundance, while the x-axis represents the m/z ratio.

**Supplementary Figure 17:**
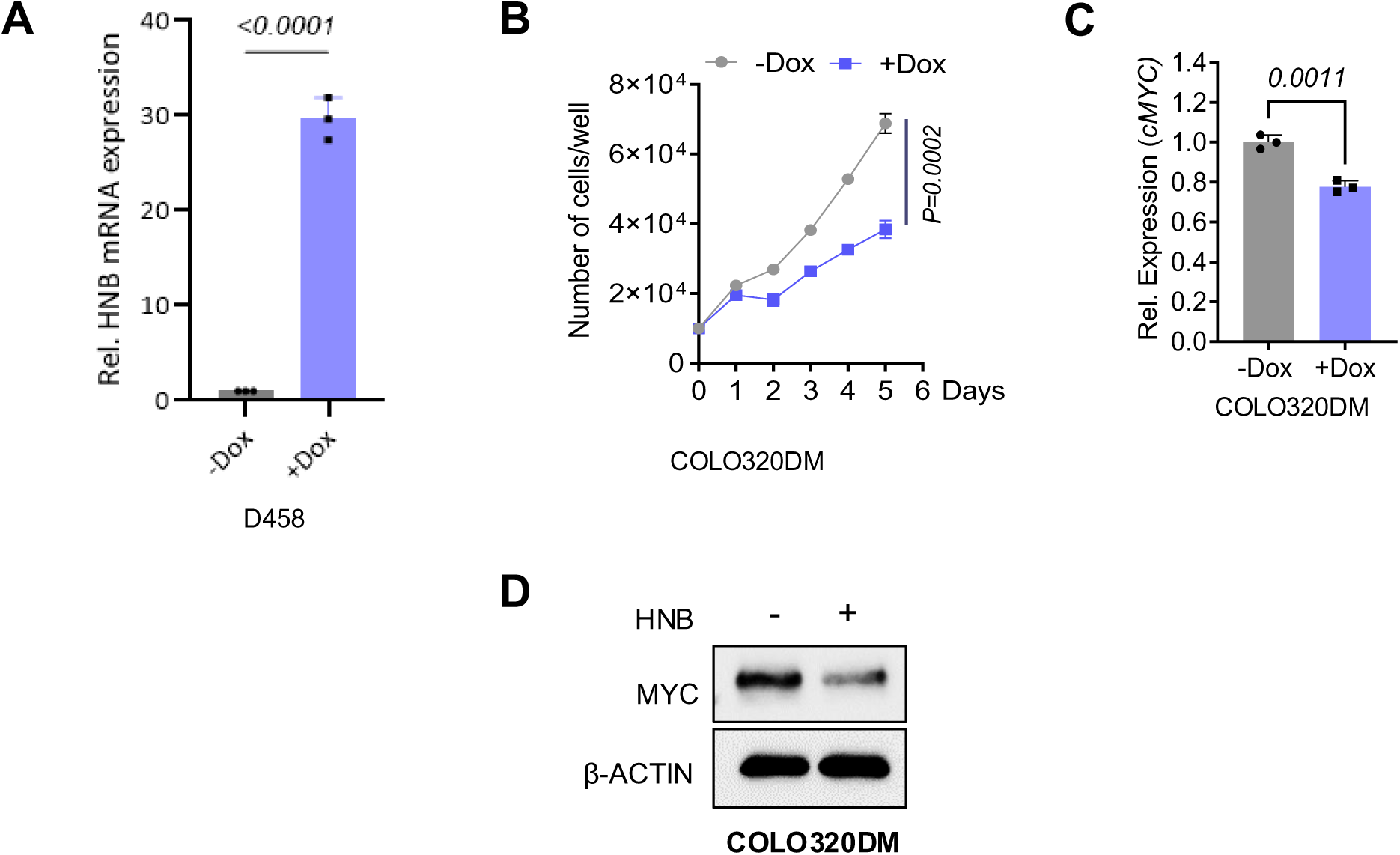
Regulation of MYC Expression by HNB: (A) Quantification of HNB transcript levels in D458 cells upon induction (+Dox), as determined by q-RT-PCR. (B) Proliferation assay of COLO320DM cells following HNB induction (-/+ Dox). (C) q-RT-PCR analysis of MYC transcript levels, with or without induction of HNB (-/+ Dox) (n=3, P values obtained through unpaired t-test). (D) Western blot analysis of MYC protein in COLO320DM cells transduced with inducible HNB transgene (-/+ Dox); β-ACTIN serves as the loading control.

**Supplementary Figure 18:**
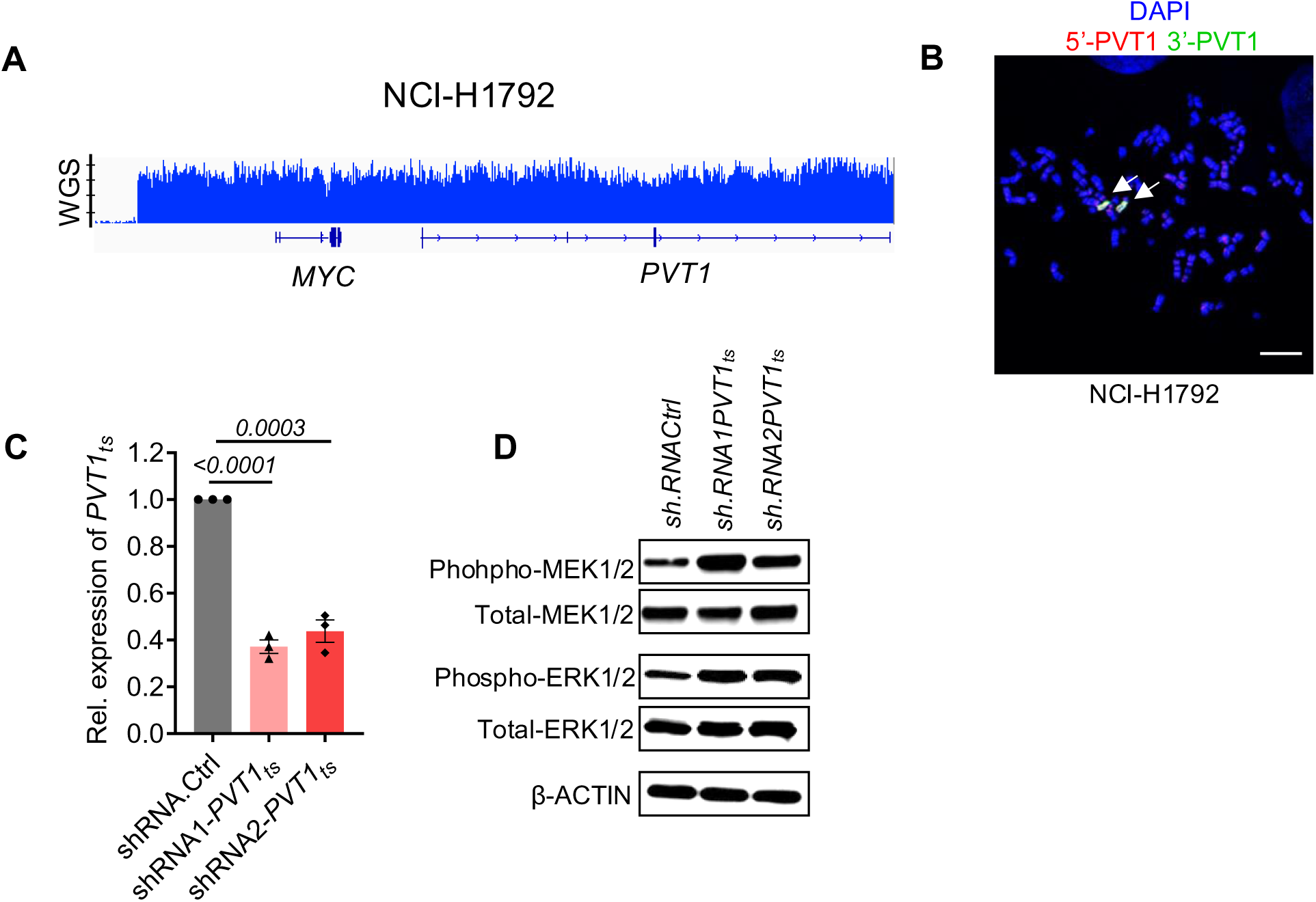
Loss of HNB activates RAS/MAPK pathway and stabilizes MYC in cell line with *MYC/PVT1* gain only (NCI-H1792) (A) Whole-Genome Sequencing (WGS) coverage across the MYC-PVT1 locus in NCI-H1792 cells **(B)** Representative Fluorescence In Situ Hybridization (FISH) using dual probes (*5’-PVT1/3’PVT1)* in NCI-H1792, revealing the intact amplified MYC/PVT1 locus (indicated by arrows). Scale bars 10 µm **(C)** qRT-PCR analysis of PVT1_ts_ transcript levels in NCI-H1792 cells transduced with sh.Cntrl, shRNA1-PVT1_ts_, and shRNA2-PVT1_ts_. **(D)** Western blot assessment of phosphorylated MEK1/2, total MEK1/2, phosphorylated ERK1/2, and total ERK1/2 in NCI-H1792 cells transduced as in (C), with β-ACTIN serving as the loading control.

**Supplementary Figure 19:**
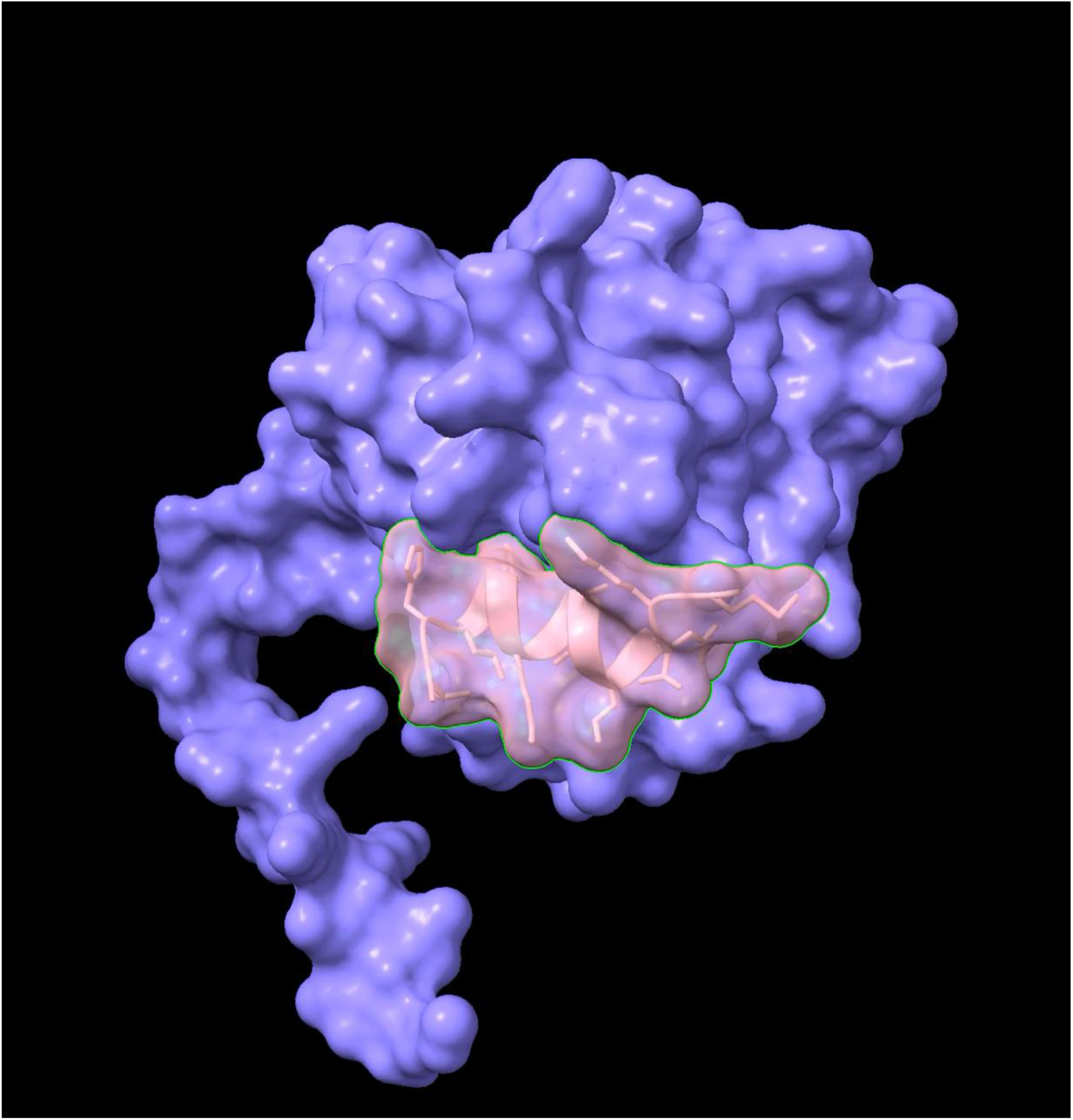
Movie showing predicted model of KRAS and HNB derived from AlphaFold 3. KRAS is represented by the purple space-filling model, while HNB is represented by the salmon ribbon and transparent space-filling models.

**Supplementary Figure 20:**
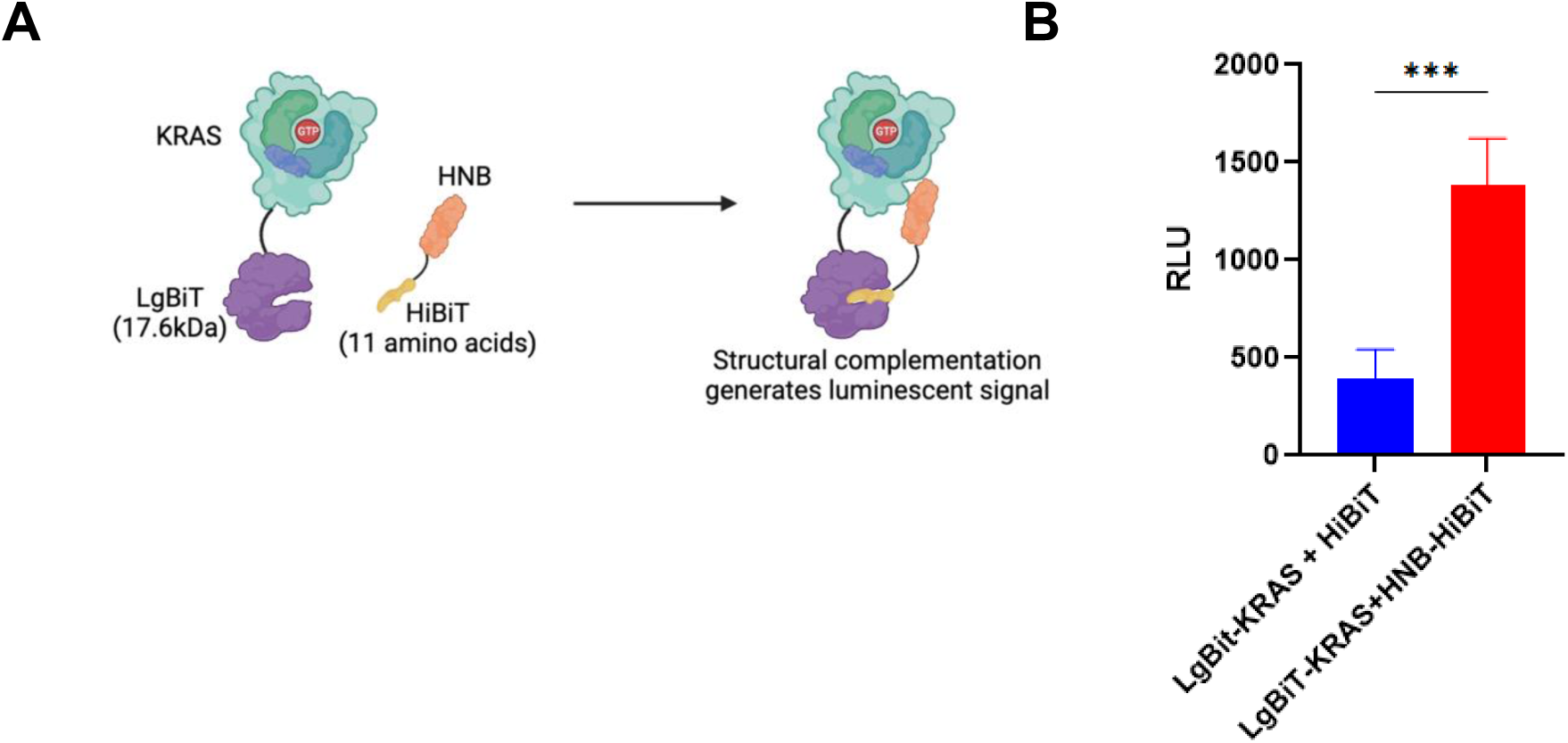
Analysis of KRAS and HNB Interaction: (A) Illustration of the NanoBiT protein-protein interaction assay highlighting the specific interaction between LgBiT-tagged KRAS and HiBiT-tagged HNB expressed in HEK293T cells. The complementation between LgBiT and HiBiT, resulting from this interaction, leads to a detectable luminescent signal. (D) Quantitative analysis of the bioluminescence arising from the LgBiT-KRAS and HNB-HiBiT interaction in HEK293T cells, with non-fused HiBiT and LgBiT KRAS serving as control. Statistical significance of the interaction is denoted (p-values obtained by unpaired t-test).

**Supplementary Figure 21:**
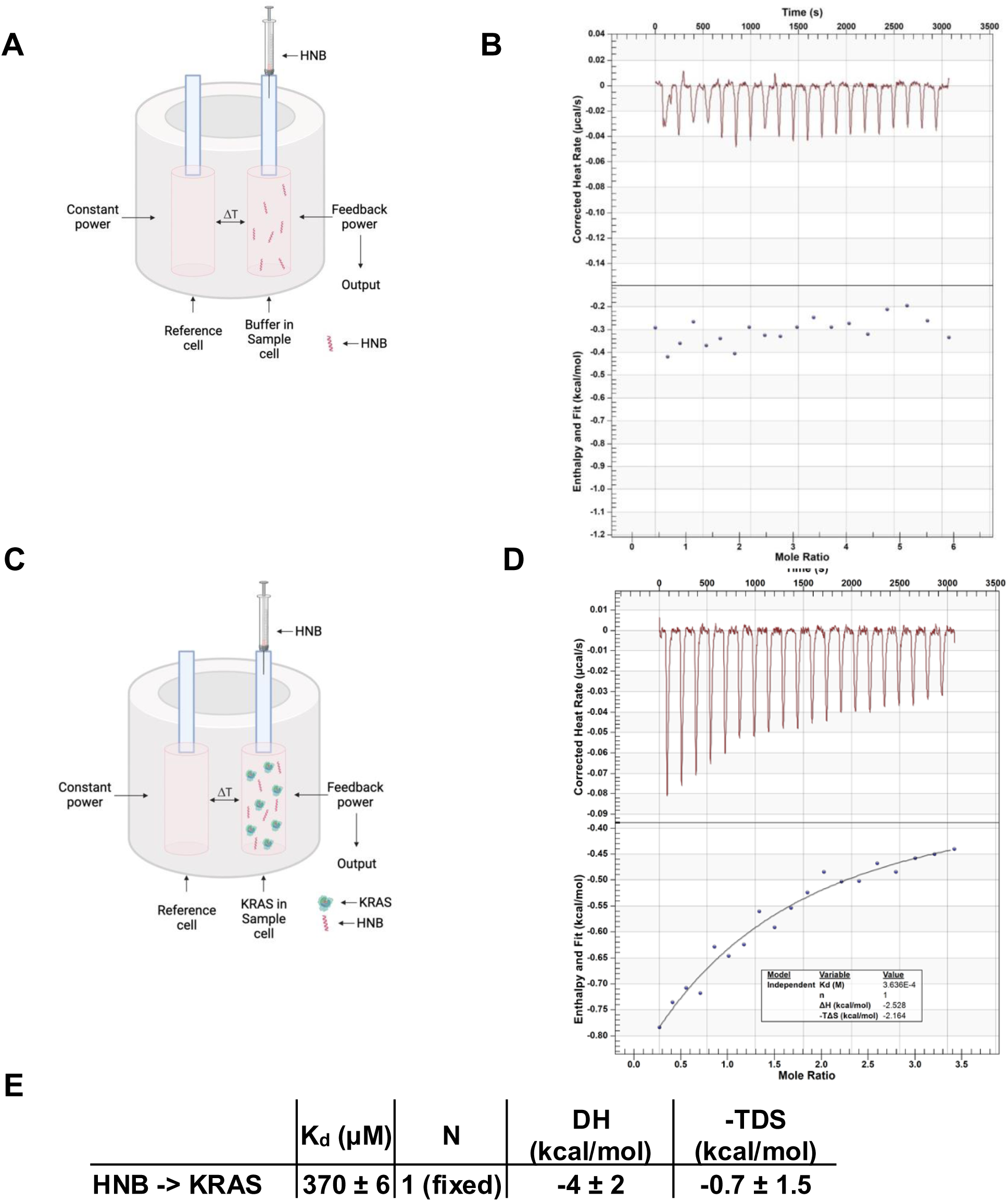
Analysis of KRAS and HNB Interaction: (A) Schematic representation of an isothermal calorimeter used for HNB titration into ITC assay buffer, detailing the injection syringe filled with HNB and the sample cell containing the ITC assay buffer. (B) Graphical display of the titration of 500 µM HNB into the ITC assay buffer, serving as a baseline control, showing the thermal response over time. (C) Schematic representation of isothermal titration calorimeter with titration of HNB with recombinant KRAS, (D,E) The plot obtained from titrating 500 µM of HNB with 93 µM recombinant KRAS in ITC buffer showing dissociation constant (Kd) of 370 ± 6 µM. Analysis was done using the NanoAnalyze software

**Supplementary Figure 22:**
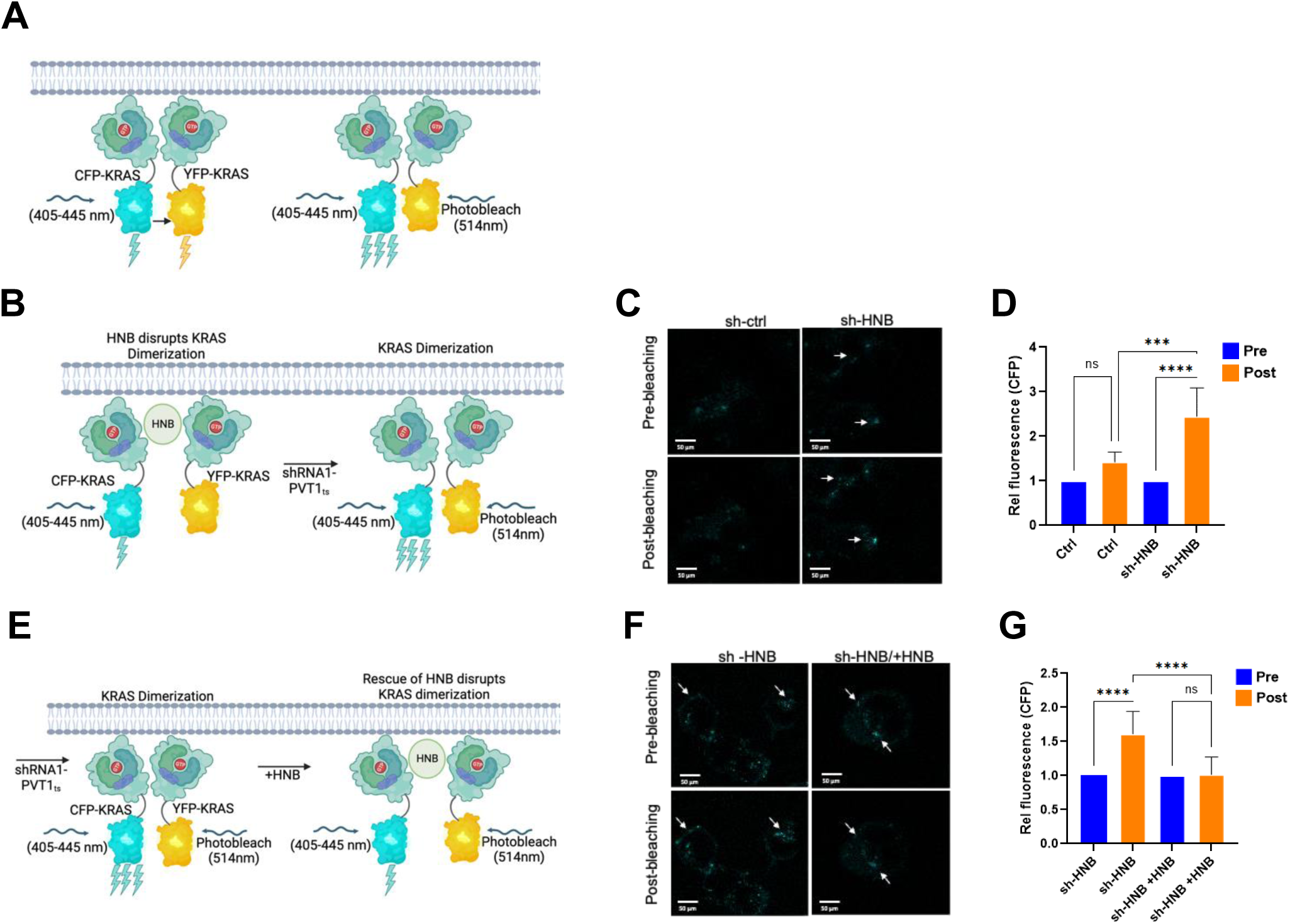
HNB regulates KRAS dimerization. (A) Schematic representation of KRAS dimerization using acceptor photobleaching FRET assay. (B) Schematic representation of KRAS dimerization on knocking down HNB using shRNA1-PVT1_ts_. (C) Representative images showing signals for CFP-KRAS and YFP-KRAS. D458 sh.Ctrl and shRNA1.PVT1_ts_ cells were co-transfected with CFP-KRAS and YFP-KRAS. Images were taken at 60X oil; the scale bar is 10μm. (D) CFP emission for KRAS in sh.Ctrl and shRNA1.PVT1_ts_ D458 cells before (Pre) and after (Post) photobleaching of acceptor. (p-value calculated by one-way ANOVA, n=10). (E) Schematic representation showing disruption of KRAS dimerization upon expression of HNB using doxycycline induction. (F) Representative images showing signals for CFP-KRAS and YFP-KRAS. D458 shRNA1.PVT1_ts_ and doxycycline-induced HNB-expressed D458 cells were co-transfected with CFP-KRAS and YFP-KRAS. Images were taken at 60X oil; the scale bar is 10μm. (G) CFP emission for KRAS in shRNA1.PVT1_ts_ and +HNB in D458 cells before (Pre) and after (Post) acceptor photobleaching. (p-value calculated by one-way ANOVA, n=10).

**Supplementary Figure 23:**
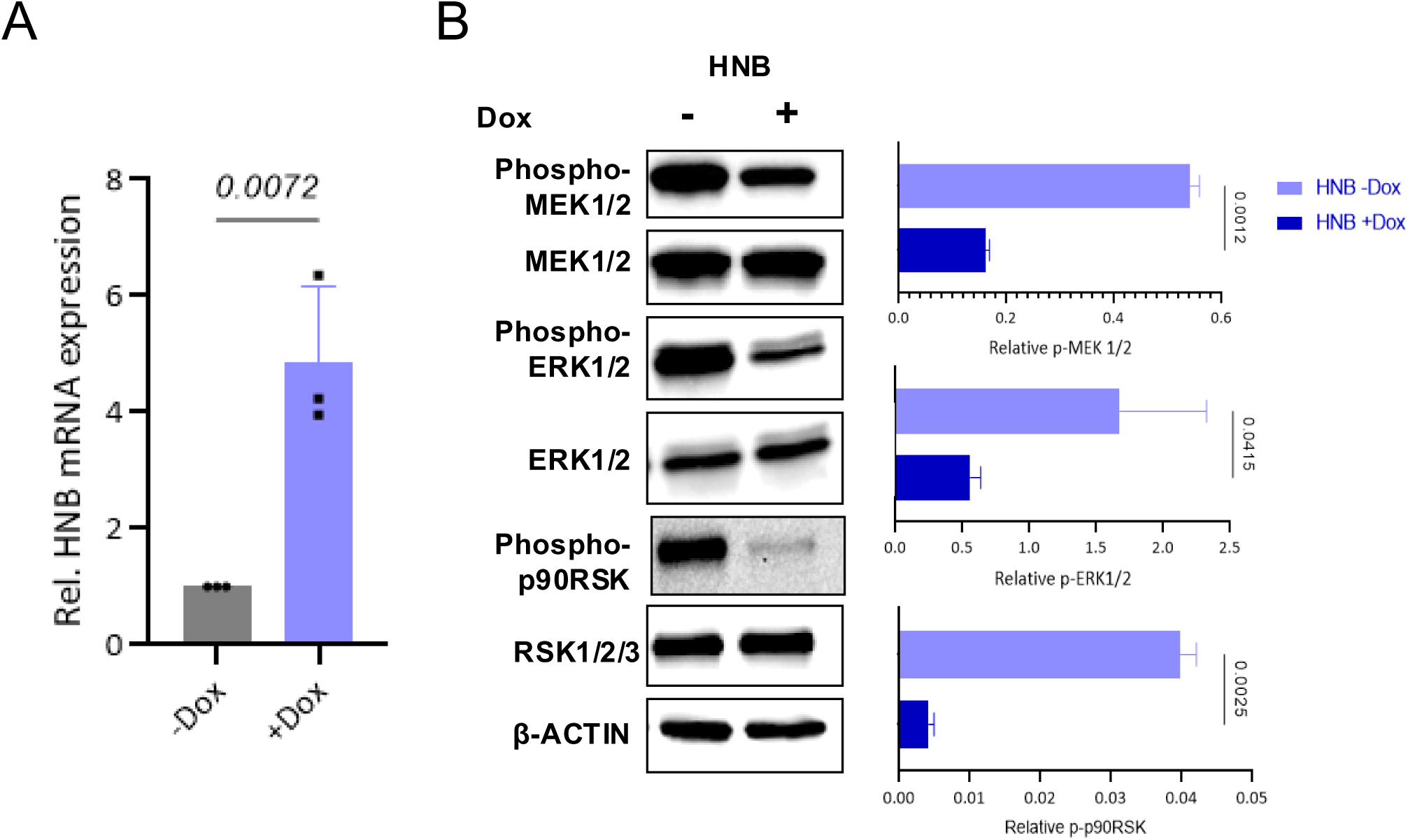
Regulation of MAPK Signaling by HNB in NCI-H1792 Cells: (A) q-RT-PCR analysis showing quantification of HNB transcript levels in NCI-H1792 cells with (+Dox) or without induction (-Dox). (B) Western blot and quantitative analysis representing the expression levels of key signaling molecules: phospho-MEK1/2, total MEK1/2, phospho-ERK1/2, total ERK1/2, phospho-p90RSK, and total RSK1-2 in NCI-H1792 cells transduced with an inducible HNB-expressing plasmid. β ACTIN is utilized as a loading control (p-values obtained by unpaired t-test).

**Supplementary Figure 24:**
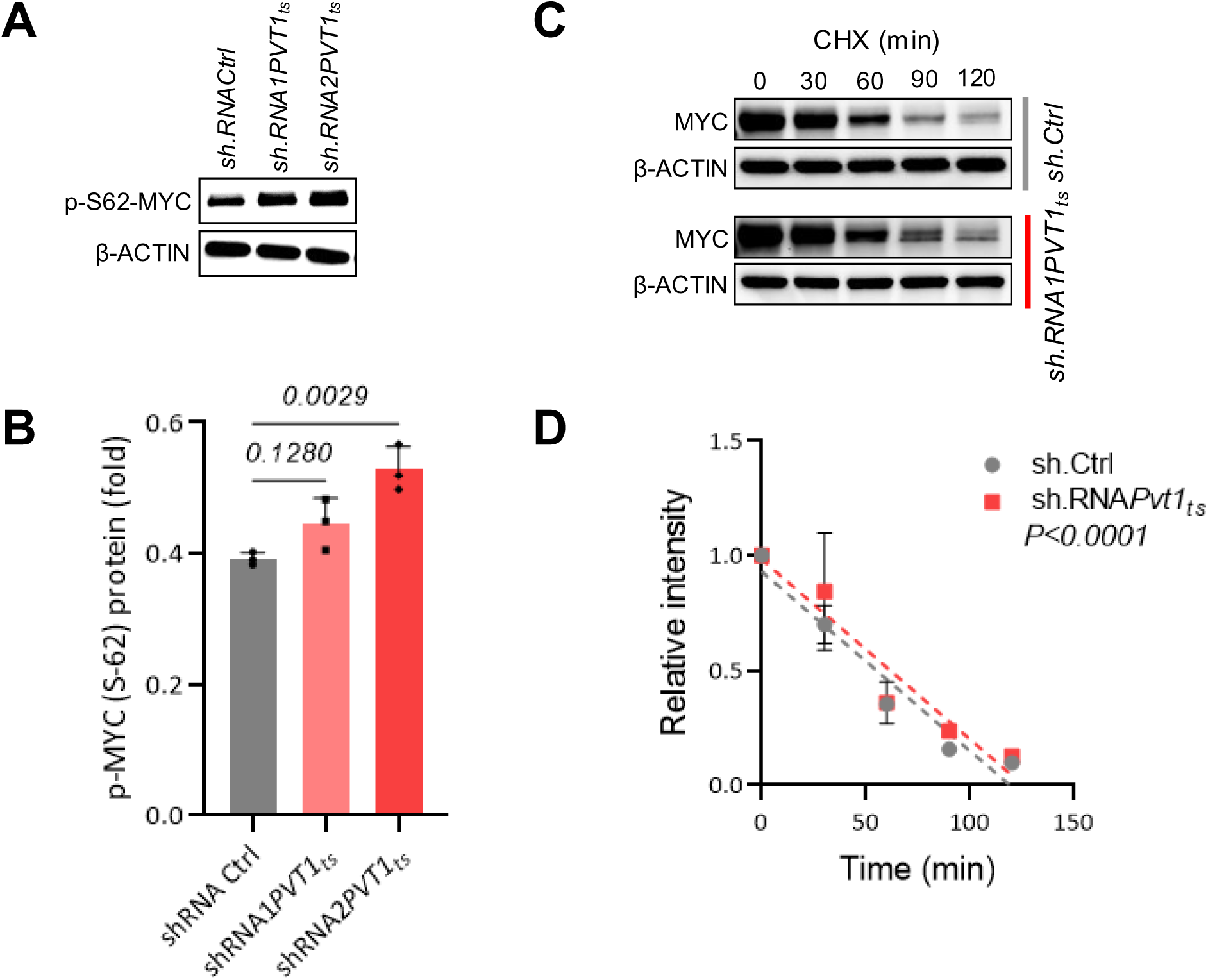
Loss of HNB enhances MYC stability in NCI-H1792 **(A,B)** Western blot and quantitative analysis of p-MYC^Ser62^ in NCI-H1792 stably transduced with sh.Cntrl, shRNA1-PVT1_ts_ and shRNA2-PVT1_ts_. β ACTIN was used as loading control ((p-values obtained by ANOVA). (C,D) Cycloheximide chase assay and quantification for MYC in NCI-H1792 cells transduced with sh.Cntrl (as control) or shRNA1-PVT1ts. β-ACTIN was used as a loading control. (P value obtained by simple linear regression)

